# Temporal evolution of color representations measured with MEG reveals a ‘coarse to fine’ dynamic

**DOI:** 10.1101/2024.12.19.629529

**Authors:** Erin Goddard, Kathy T. Mullen

## Abstract

Color perception is based on the differential spectral responses of the L, M and S-cones, subsequent subcortical and cortical computations, and may include the influence of higher order factors such as language. Although the early subcortical stages of color vision are well characterised, the organization of cortical representations of color remain elusive, despite numerous models based on discrimination thresholds, appearance and categorization. An underexplored aspect of cortical color representations is their dynamic evolution. Here we compare the evolution of three different color representations over time using magnetoencephalography (MEG). We measured neural responses to 14 hues at each of 3 achromatic offsets (increment, isoluminant and decrement) while participants attended either to the exact color of the stimulus or its color category. We used a series of classification analyses, combined with multidimensional scaling (MDS) and Representational Similarity Analysis, to ask how cortical representations of color unfold over time from stimulus onset. We compared the performance of ‘higher order’ models based on hue and color category with a model based simply on stimulus cone contrast and found that all models had significant correlations with the data. However, the unique variance accounted for by each model revealed a dynamic change in hue responses over time, that was consistent with a ‘coarse to fine’ transition from a broad clustering into categorical groups to a finer within category representation. Notably, these dynamics were replicated across datasets from both tasks, suggesting they reflect a robust reorganization of cortical hue responses over time.

## Introduction

The first stage of human color vision is based on the combination of the differential responses of the L, M and S-cones to the visual stimulus. In this first postreceptoral stage, comparisons of the cone responses are initially made within two distinct cone opponent streams, an L/M cone opponent response (loosely called ‘red-green’) and one based on the opponent combination of S-cones with the other two (loosely called ‘blue-yellow’). Based on physiological and behavioural data, these two pathways are thought to remain segregated within the subcortical processing stages, using different anatomical cell types and pathways in the retina, LGN, and with distinct V1 cortical inputs (Derrington et al., 1984; Lennie & Movshon, 2005). There is also evidence for a behavioural separation at detection threshold (Eskew, 2009; Mullen & Sankeralli, 1999). These two low-level pathways, however, do not provide adequate accounts of color appearance or the perceptual organization of the different colors. For example, colors that are equally spaced in a representation based on perceptual differences/similarity (e.g. using a CIE L*a*b* space), vary in a non-uniform way in a space based on the relative cone responses (e.g. a cone contrast space). Furthermore, it has been widely observed that stimulating the cardinal directions of a cone contrast space, which uniquely activate one type of cone opponent process at a time, do not generate the color appearance of a unique hue. Hence, the unique hues are not aligned with the cone opponent axes and further computations are required for cone opponency to generate the full range of color appearances (Conway et al., 2023; de Valois & de Valois, 1993; Li et al., 2022; Stockman & Brainard, 2010). Embedded within this perceptual color space is a categorical organization, in which colors can be placed into groups according to their color appearance, such as the 11 basic color names of Berlin and Kay (1969). Whether there are any categorical effects in color perception, including potential interaction with language, remains controversial, but evidence is clearer for ‘categorical facilitation’ effects, likely mediated by a shift of attention to the linguistic distinction between color categories (for a review, see Witzel, 2019). So far, the computational nature of the processes underlying categorical color grouping and subsequent color naming also remains unknown (Siuda-Krzywicka et al., 2019).

Imaging approaches provide an important route into the understanding of how colors are represented in the cortex. fMRI can be used to decode stimulus color using a multivariate approach to analyse spatially distributed patterns of voxel activity in cortex (Brouwer & Heeger, 2009, 2013; Goddard et al., 2010; Goddard & Mullen, 2020; Parkes et al., 2009). Although colors can be decoded in early visual cortical areas (V1, V2, V3, hV4 & VO), the ventral areas hV4 and VO showed a stronger perceptual relationship than earlier areas such as V1 and V2, with perceptually similar colors evoking the most similar responses (Brouwer & Heeger, 2009). Brouwer & Heeger (2013) also found that areas hV4 and VO1 showed evidence of categorical clustering, especially when participants were engaged in a color categorization task, compared to a diverted attention condition. There is also evidence of further areas of high color responsiveness more anterior in the ventral visual cortex (Komatsu et al., 1992; Stoughton & Conway, 2008; Zeki & Bartels, 1999), including adjacent to areas associated with higher-order responses such as to objects, faces and places (Lafer-Sousa et al., 2016; Lafer-Sousa & Conway, 2013).

MEG signals can be used to decode stimulus color with the advantage of allowing the precise timing of the stimulus response to be determined (Hermann et al., 2022; Rosenthal et al., 2021). This previous work, using 4 stimulus colors and two luminance levels, has shown that MEG signals can be used to decode stimulus color for over 300ms after stimulus onset (Rosenthal et al., 2021), and provides a proof of concept that similarity relationships can be used to test the geometry of stimulus color representations. Other MEG studies have also demonstrated color decoding from MEG signals (Goddard, Carlson, et al., 2022; Goddard, Shooner, et al., 2022; Sandhaeger et al., 2019; Teichmann et al., 2019, 2020), and task-based changes in color coding consistent with sharpening of color tuning with attention (Bartsch et al., 2017; Goddard, Carlson, et al., 2022). In addition, there are many examples of decoding stimulus color from EEG signals (Bae & Chen, 2024; Chauhan et al., 2023; Grootswagers et al., 2024; Hajonides et al., 2021; Retter et al., 2023; Rozman et al., 2024; Sutterer et al., 2021; Wu et al., 2022).

Here we recorded MEG signals while participants viewed a set of 42 stimuli that varied in hue as well as achromatic offset and compared the timing of the emergence of neural representations of chromatic and achromatic information. We included two task conditions, both involving attention to the stimulus color but only one requiring color naming (categorization), to test whether the task of color naming influences the way color is represented in the cortex.

To test how neural color representations evolve over time, we first used an exploratory analysis of the MEG data (multi-dimensional scaling, MDS), and second, a model-driven Representational Similarity Analysis (RSA) (Kriegeskorte, Mur, & Bandettini, 2008; Kriegeskorte, Mur, Ruff, et al., 2008; Popal et al., 2019). In the RSA, we compared three different theoretical (model) response spaces, reflecting different factors that could be influential in the neural representation of color: the cone contrast differences between the stimuli; the perceptual differences between stimuli based on color discriminability; and color category, in which within-category pairs of stimuli are predicted to produce more similar responses than cross-category ones. Our results show an orderly representation of hue across the stimulus-induced response that is broadly consistent with each of the three *a priori* models above. Interestingly, we find a non-linear warping of the representational space that is dynamic, changing over time from the stimulus onset. These nonlinearities suggest a ‘coarse to fine’ dynamic, in which the cortical hue representation includes an initial clustering of similar hues before a later separation of these clusters to represent the finer differences between stimuli. We replicated the dynamics of these hue representations across the data from the two different tasks, and finding only a subtle indication of possible task-mediated differences in the later part of the response.

## Materials and methods

### Participants

We collected MEG data on 8 participants (5 female, 3 male, aged 21-34 years, mean=26.4 years). Each participant had no history of neurological and/or psychiatric disorders, normal or corrected-to-normal visual acuity, and normal color vision as assessed with Ishihara plates (Ishihara, 1990) and the Farnsworth-Munsell 100-hue test (Farnsworth, 1957). All participants provided informed consent prior to taking part, and all experimental procedures were approved by the Ethics Review Board of the McGill University Health Centre and were conducted in accordance with the Declaration of Helsinki. We collected MEG data for each participant during a single session, of approximately 2 hours. Each participant was familiarised with all tasks prior to the day of the MEG data collection, but all behavioural data reported here are from the MEG session.

Our sample size (n=8) is smaller than previous work testing for color representations with MEG (n=18, Hermann et al., 2022; Rosenthal et al., 2021), but larger than earlier work using fMRI (n=5, Brouwer & Heeger, 2009, 2013), and larger than our recent MEG work which included similar analysis approaches (n=6, Goddard, Shooner, et al., 2022). Teichmann et al. (2022) used real and simulated data to test the effect of participant and trial numbers on statistical power for classification analyses of MEG responses to visual stimuli. They report that sufficient data at a subject level (i.e. enough trials) is likely more important for statistical power than sample size, giving the empirical example of a data set with 9 participants completing 1600 trials having equivalent power to data from 18 participants who completed 400 trials. In this study, we prioritised many trials (>3,800 trials per participant) to increase this subject level power.

### Visual stimuli

All visual stimuli were large circular patches of uniform color, serially presented. Each circular stimulus was 40 degrees in diameter, presented in a raised-cosine envelope, and outside the stimulus area the screen was set to a grey of mean luminance, as illustrated in **Figure 1**. We used 14 hues, chosen to be equally spaced around a circle of diameter 40 in the isoluminant plane of CIE Lab space. The saturation of the stimuli was chosen to be close to the maximum achievable within the gamut of the display.

**Figure 1:**
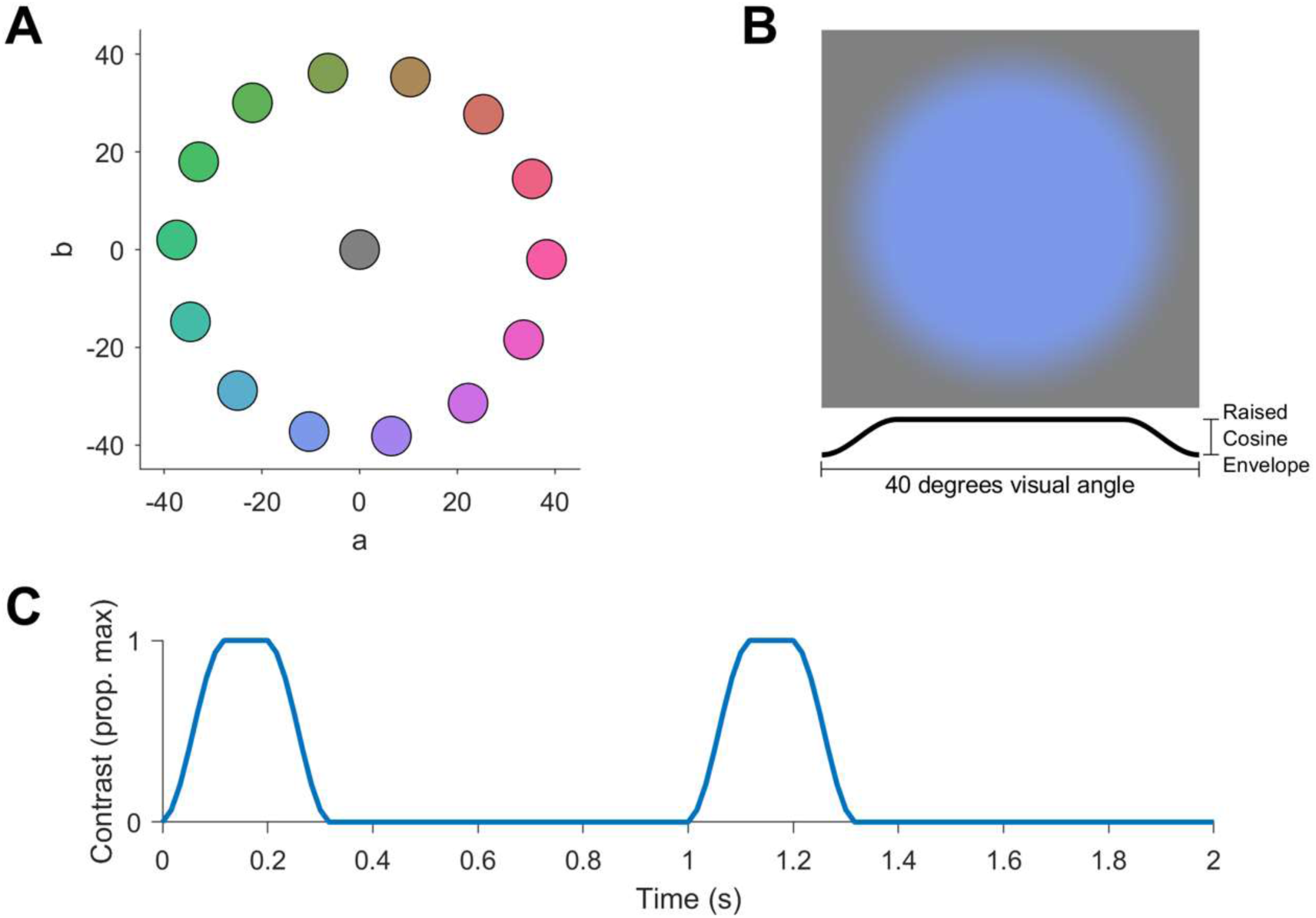
Spatial and temporal stimulus properties. **A**: Stimuli were 14 hues, equally spaced around a circle in the isoluminant plane of Lab space, presented at 3 luminance levels (not shown here). The background was held at mean grey (a=b=0). **B**: Each stimulus was a color presented in a circular raised-cosine spatial envelope. **C:** Example timecourse of 2 trials: each stimulus was presented in a temporal raised-cosine envelope.

We determined the isoluminant plane of CIE Lab space for each subject individually using heterochromatic flicker photometry (He et al., 2020; Smith & Pokorny, 1975). At the start of the experimental session, participants viewed a sinusoidally flickering stimulus, alternating between a frame of a colored stimulus (as in **Figure 1B**), and an achromatic stimulus of the same spatial properties. Participants used a mouse to adjust the luminance of the achromatic stimulus until their impression of flicker was minimised, indicating their selection with a mouse click. After performing at 3 settings for each of the 14 hues, we averaged the selected luminances for each hue. We fit a sinusoid to these average offsets across hue direction, with amplitude and phase as fitted parameters, and used the fitted values to adjust the experimental stimuli for perceptual isoluminance. Across participants, the phase of the fitted sinusoid aligned with stimulus hue such that the maximum correction was applied to the stimuli closest to the L-M isolating axis, as expected for a correction to adjust for individual variability in the luminance mechanism. The maximum amplitude of this achromatic correction (L=M=S) varied across individuals (range 0.8-4.9%, mean 2.9% cone contrast).In addition to the 14 hues that were isoluminant with the background, we used another 28 hues that were of the same hue direction but with a 12.5% luminance increment or decrement, giving a total of 42 unique stimuli.

After the stimuli were calibrated for their individual perceptual isoluminance, participants viewed all stimuli sequentially, and verbally reported the color category of each stimulus, which were recorded by the experimenter. Participants were restricted to using the labels ‘red/pink’, ‘orange’, ‘yellow’, ‘green’, ‘blue’ and ‘purple’. Each stimulus was presented 3 times, totalling 126 trials. The modal color labels for each unique stimulus were used to define each participant’s color categories, used in the category task (described under ‘participant task’ below) and the category model (used in the Representational Similarity Analysis, described below).

No MEG data were collected during these preliminary tasks.

### MEG methods: acquisition protocols

MEG data were collected with a whole-head MEG system (CTF OMEGA System) consisting of 275 axial gradiometers. For each MEG session we first collected 5 minutes of empty room recordings which we used to estimate noise covariance of the sensors (see below). Prior to the participant entering the magnetically shielded room, three marker coils were placed on the participant’s head. Marker positions, nasion, left and right pre-auricular points, and the participant’s head shape were recorded with a pen digitizer (Polhemus Isotrak, Colchester, VT), using a minimum of 500 points. Two EOG electrodes were places above and below the left eye, to record eye blinks and eye movements during the MEG session. Two electrodes were placed across the plane of the chest to collect electrocardiographic (ECG) signals, and a reference electrode was placed below the participant’s collarbone. Each participant’s MEG data and simultaneous EOG and ECG signals were collected at a sampling frequency of 2400Hz. In conjunction with these data, we collected participants button responses using a VPixx ResponsePixx button box system (VPixx Technologies, Canada).

### MEG methods: display apparatus and calibrations

We displayed stimuli using a PROPixx DLP LED projector (VPixx Technologies, Canada, resolution 1920×1080), located outside the magnetically shielded room, to back-project images onto a custom screen via two mirrors. Participants, lying supine in the MEG system, viewed the custom screen, located above them, from a distance of 45cm. We used a Windows PC (Windows 7) running MATLAB (R2017a) in conjunction with routines from Psychtoolbox 3.0 (Brainard, 1997; Kleiner et al., 2007; Pelli, 1997) to generate and project the stimuli (refresh rate 60Hz, mean luminance 106 cd/m2). The PROPixx DLP LED projector has a linear gamma, and was color calibrated as described previously (Michna et al., 2007; Mullen et al., 2007, 2008). We precisely aligned stimulus presentation times with the recorded MEG data using the VPixx ‘Pixel Mode’ to record the output of a single pixel along with the MEG data.

### MEG methods: experimental design and participant’s task

Each participant completed the MEG data collection in a single session, which was divided into ten blocks of 5-7 minutes each, with breaks between blocks. On each block of trials, participants saw the same stimulus set (summarised in Figure 1) and performed a 1-back task requiring color categorization or discrimination. In both tasks, the spatial and temporal properties of the stimuli were identical, with each trial comprising a single color presented in a temporally raised-cosine envelope (300ms in total, with 100ms onset, 100ms sustained presentation, and 100ms offset). Across both tasks, the trials were presented in a continuous stream, and participants responded with a keypress whenever 2 consecutive stimuli were an exact match (discrimination task), or a color category match (category task, where same category was defined as stimuli that were given the same label in the color naming task).

For both tasks, we used at least 42 trials of each of the 42 unique stimuli (1764 trials). These were presented in a counterbalanced order where each stimulus was preceded by each of 42 stimuli at least once. Since category matches were more common than exact stimulus matches, we added extra exact match trials to the discrimination task blocks so that the average rate of match trials was the same across the two tasks. The total number of trials in the discrimination task ranged from 2066 to 2130 across participants, since the match rate in the category task varied with their individual color labels.

In pilot testing, participants were faster and more accurate for the discrimination task than for the category task: to approximately equate the tasks for difficulty in the experiment we used a longer inter-stimulus interval (ISI) in the category task (700ms) than in the discrimination task (450ms), giving stimulus onsets that were separated by 1000ms and 750ms respectively.

For both the category and the discrimination tasks we evaluated participants’ performance using their hit rate (HR: proportion of match trials with a button press) and false alarm rate (FA: proportion of non-match trials with a button press) to calculate sensitivity (d’) using the MATLAB function *norminv* (inverse of the cumulative normal distribution):

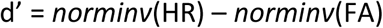

All participants performed these tasks well above the chance rate of d’=0. Across participants, the average d’ = 2.7 (range d’=1.8-4.1) on the category task, and d’ = 3.3 (range d’=2.4-4.2) on the discrimination task.

### MRI methods: retinotopic and functional localisers

Each participant completed an MRI session in which we acquired high-resolution anatomical images of their brains and functional data used to define regions of interest. All magnetic resonance imaging took place at the McConnell Brain Imaging Centre, McGill University, Montreal, Canada. For each participant we acquired two high-resolution three-dimensional whole head T1 images using an MP-RAGE sequence (TI=900ms, TR = 2300ms, TE = 3.41ms, 1.0mm^3^ resolution), and averaged these two images to generate the participant’s anatomical template. Functional T2* MR images were acquired on a 3T Siemens MAGNETOM Prisma system with 32-channel head coil. Gradient-echo pulse sequences were used to measure blood oxygenation level-dependent (BOLD) signal as a function of time. We identified the visual cortical regions V1, V2, V3, V3A/B, LO1/LO2 and hV4 for each participant using rotating wedge stimuli and expanding and contracting concentric rings (Engel et al., 1994; Sereno et al., 1995), standard definitions of these areas (Brewer et al., 2005; Goddard et al., 2011; Larsson & Heeger, 2006), and the foveal confluence (Schira et al., 2009). To localize areas VO1, VO2 and hMT+ we used data from the retinotopic mapping scans in conjunction with functional localisers for VO (Mullen et al., 2007) and hMT+ (Huk et al., 2002). Full details of our retinotopic mapping procedures, including scanning protocols, data preprocessing and area definition have been described previously (Goddard et al., 2019).

### MEG data analysis: preprocessing and source reconstruction

Preprocessing, forward modeling and source reconstruction of MEG data were performed using Brainstorm (Tadel et al., 2011, http://neuroimage.usc.edu/brainstorm). For each participant’s template anatomical (a high-resolution MRI image), we used the automatic segmentation processes from Freesurfer 6.0 (Dale et al., 1999; Fischl et al., 1999) to define the grey/white matter and pial/grey matter boundaries. Using Brainstorm, we imported the output of Freesurfer and created a 15,000-vertex model of each participant’s cortical surface. For each block of trials in the MEG data, we aligned the participant’s cortical surface model to the median measured marker coil locations for that block by aligning the head shape data from the MRI with the head shape relative to the marker coils, as recorded with the pen digitiser. For each functional run, we generated a forward model for each model by applying a multiple spheres model (Huang et al., 1999) to the participant’s cortical surface model at this measured head location.

Functional data were preprocessed in Brainstorm with notch filtering (60, 120 and 180Hz), followed by bandpass filtering (0.2-200Hz, using the Brainstorm default of an even-order linear phase FIR filter). We preprocessed data from the empty room recording using identical protocols, then used the output to estimate the noise covariance for the session. Cardiac and eye blink artifacts were removed from functional data using signal space projection (SSP): cardiac and eye blinks events were identified using default filters in Brainstorm, manually verified, then used to estimate a small number of basis functions corresponding to these noise components, which were removed from the recordings (Uusitalo & Ilmoniemi, 1997). From these functional data we extracted an epoch of data for each trial: from -100 to 1000ms relative to stimulus onset, and downsampled to 200Hz. Using the noise covariance estimate, regularized using the median eigenvalue, we applied a minimum norm source reconstruction to this trial data.

For classification analyses we generated three datasets using each participant’s functionally defined cortical areas, as in previous work (Goddard, Shooner, et al., 2022). In the ‘early visual cortex’ (EVC) region of interest, we included data from all vertices located within areas V1, V2 and V3, while the ‘ventral visual cortex’ (VVC) region of interest included areas hV4, VO1 and VO2, and ‘dorsolateral visual cortex’ (DVC) included areas V3A/B, LO1, LO2 and hMT+. We found very similar effects across these ROIs, which we believe most likely reflects the failure of the source reconstruction to completely isolate responses from these adjoining cortical regions. Since there is little difference across ROIs, below we show data from EVC only, but equivalent figures for VVC and DVC are included in the Appendix.

### MEG data analysis: classification-based analyses

We used a series of classification analyses to measure the amount of information about stimulus color and luminance that was available in the neural responses, as measured with MEG. We repeated these analyses for each 5ms bin to capture how this information changed over time. Out of the total 15,000 sources in each participant’s head model, the EVC ROI included an average of 731 sources (*SD* = 165), the VVC ROI included an average of 396 sources (*SD* = 67), and the DVC ROI included an average of 434 sources (*SD* = 106). We reduced these with PCA and retained data from the first *n* components, which accounted for 99.9% of the variance, for the classification analysis (mean = 285.9, *SD* = 13.7 for EVC; mean = 283.6, *SD* = 17.4 for VVC; mean = 283.6, *SD* = 20.5 for DVC).

We conducted a series of pairwise classifications based on either stimulus hue or stimulus luminance, with separate analyses for data from the category and discrimination tasks, and for each ROI. In each case, we trained classifiers to discriminate between two categories of trial (i.e. two hues or two luminance levels) and tested on held-out data. We report results obtained with a linear support vector machine (SVM) classifier, using the MATLAB function *fitcsvm* with ‘*KernelFunction’* set to *‘linear’*. For all analyses we expressed average classifier accuracy in *d’* (a unit-free measure of sensitivity). Chance classification performance yields *d’* = 0.

For classification analyses, we generated ‘pseudo-trials’ to increase signal-to-noise ratio along the dimension of interest, and to reduce data variability due to stimulus dimensions orthogonal to the dimension of interest (e.g. Goddard & Mullen, 2021). When training classifiers to discriminate stimulus hue, we used pseudo-trials that were each the average of 12 trials of the same hue: 4 of each luminance level. Trials were sampled without replacement, to ensure that each pseudo-trial was comprised of independent data. For data from a single task, this yielded 11 pseudo-trials for each of the 14 hues. For each pair of hues, we used these pseudo-trials in an 11-fold cross-validation, in each case training on data from 20 pseudo-trials (10 from each hue) then testing the classification rule on its classification on the held-out pair of pseudo-trials. We repeated this entire process 10 times, with different partitions of the data into pseudo-trials. We performed an equivalent procedure when classifying stimulus luminance: generating pseudo-trials that were each the average of 14 trials of the same luminance, 1 of each stimulus hue. This yielded 42 pseudo-trials for each of the 3 luminance levels, and for each pair of luminance levels we used a 42-fold cross-validation.

For both hue and luminance, we performed these classification analyses on data from within each 5ms time bin, to see how information about these stimulus features in the neural responses unfolded over time. For decoding of stimulus hue in the EVC ROI, we additionally tested for cross-temporal generalization: training classifiers on data from one time bin and then testing this classification rule on data from each later time bin in held-out pseudo-trials.

### MEG data analysis: Representational Similarity Analysis (RSA) and Multi-dimensional Scaling (MDS)

When decoding stimulus hue, pairwise comparisons of the 14 unique hue directions resulted in a 14×14 dissimilarity matrix (DSM) for each time bin where each cell of the matrix was defined by classifier accuracy. To visualise how the neural representations of color were unfolding over time, we used multidimensional scaling (MDS) to reduce these matrices to the best-fitting two-dimensional representation of the 14 hues. Each MDS was completed using the MATLAB function *mdscale* with ‘*criterion’* set to *‘metricstress’* (minimising the stress normalised with the sum of squares of the dissimilarities).

We also used Representational Similarity Analysis (Kriegeskorte, Mur, Ruff, et al., 2008; Nili et al., 2014) to compare these matrices with three *a priori* models, based on different representations of stimulus hue (**Figure 2**). Each of these models predicted that the dissimilarity of the neural responses would be proportional to the stimulus difference, but with stimulus difference quantified in three possible ways: as Euclidian distance between stimuli in cone contrast (Figure 2A), CIE L*a*b* space (Figure 2B), or in color category (Figure 2C). For each time bin, we rank-correlated each participants’ observed DSM with each of these three models. We compared model performance against the benchmark of the ‘noise ceiling’, which estimates the expected correlation of the observed data with a hypothetical (unknown) ‘true’ model, given the noise in the data (Nili et al., 2014). The upper-bound of the noise ceiling is calculated by correlating the group average DSM at each time bin with each individual’s DSM, which, due to overfitting, will overestimate the true model’s correlation. The lower-bound of the noise ceiling is calculated by correlating each individual’s data with a group average that excludes their own data, avoiding overfitting, and providing a lower bound on the expected correlation with the true model.

**Figure 2:**
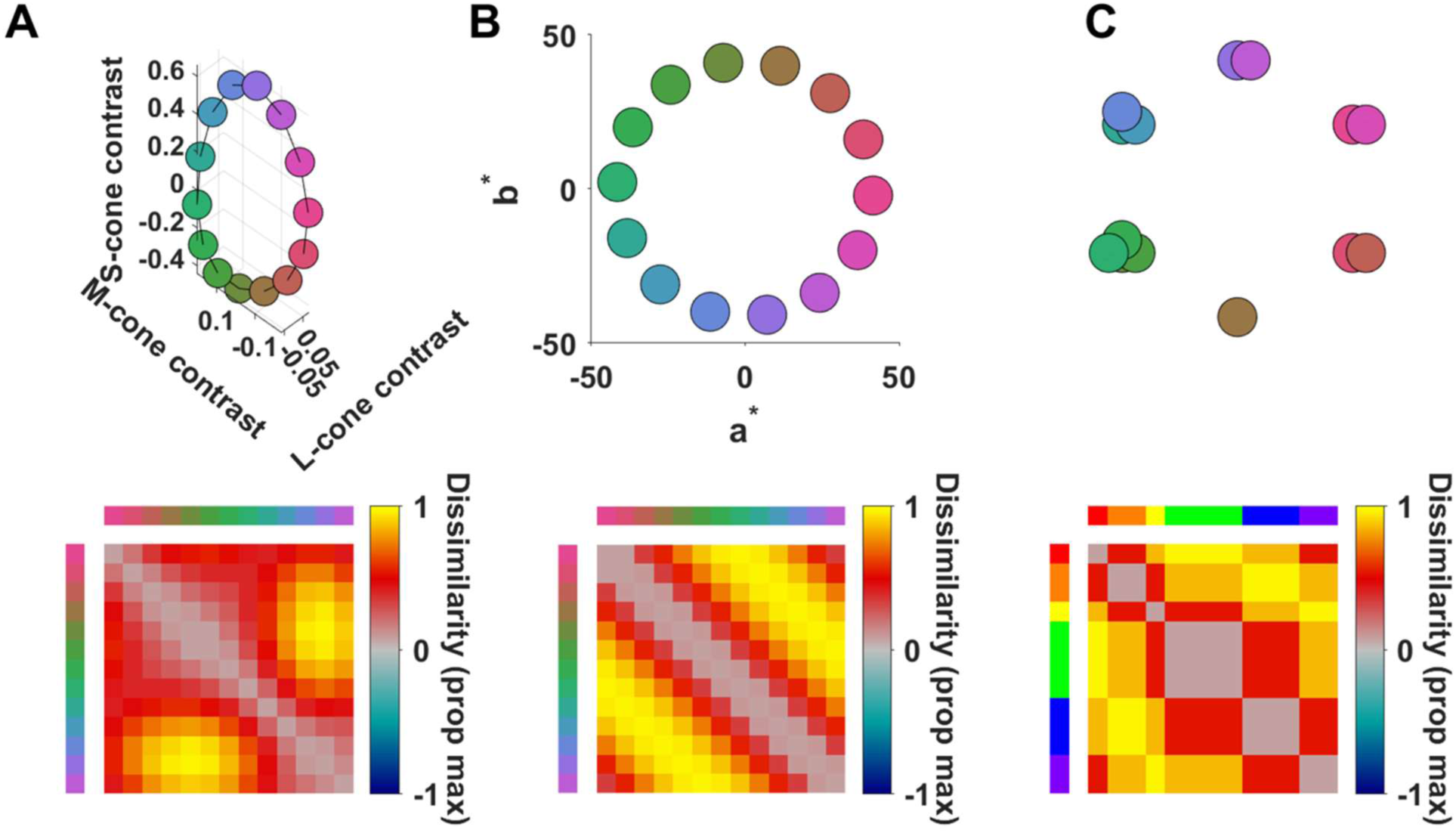
Model dissimilarity matrices (DSMs) based on hue for Representational Similarity Analysis (RSA). We tested classifier data against 3 models of dissimilarity for the 14 stimulus hues, based on stimulus cone-contrast (**A**), hue direction in CIE L*a*b* space (**B**) or color category (**C**). In each case, the upper plot indicates the stimulus representation according to the relevant model, and the low plot shows the corresponding model DSM. The hues were selected to be equally spaced in CIE L*a*b* space (**B**); when replotted in a cone contrast space (**A**), the hues are not equally spaced, but elongated along the S-cone contrast axis, while the category model (**C**) treats within-category hues as identical, grouped into 6 clusters equally spaced around a circle. The category model shown above (**C**) is an example model for one participant; in the analysis, each participant’s color naming data were used to generate an individual category model. These three models were positively correlated with one another, Spearman’s rho r = 0.72 (cone contrast vs hue), r = 0.25-0.44 (cone-contrast vs category), r = 0.68-0.73 (hue vs category), with variation in correlation with the category model due to inter-participant variation in color naming.

Since our *a priori* models are positively correlated with each other (see legend of Figure 2 for correlation values), they will tend to co-vary in their correlation with the data. To estimate the extent to which these models predicted unique variance in classifier performance, we fit a series of general linear models (GLMs). For each GLM, we used the 3 models as regressors, applying *spm_orth.m* (from SPM12, www.fil.ion.ucl.ac.uk/spm/doc) to perform recursive Gram-Schmidt orthogonalization (Golub & Loan, 1996) of these regressors so that the final regressor would only capture variance that is predicted by the final model and orthogonal to the other two models. We then converted each data matrix to ranked classification accuracy values and fit these ranks with a GLM comprised of the orthogonalized regressors, using the MATLAB function *fitglm.m*. We repeated this process three times, so that each model was the final regressor in one GLM, providing the parameter estimates (beta weights) which indicated any unique variance captured by this model.

When decoding stimulus luminance, the pairwise comparisons resulted in a 3×3 DSM. We conducted an RSA on these data, testing the data on two models, based either on luminance intensity or luminance contrast. Both models are shown in insets in Figure 5.

### Statistical analyses

To test whether classifier accuracy was above chance, and to test whether model correlations and parameter estimates (beta weights) were above zero, we used Bayes Factor analyses, an alternative to the traditional frequentist approach (Kass & Raftery, 1995; Morey & Wagenmakers, 2014). A Bayes Factor compares evidence for competing hypotheses; here we report where there is moderate (BF>3) or strong (BF>10) evidence in favour of the alternate hypothesis, or at least moderate (BF<1/3) evidence in favour of the null hypothesis. We implemented all Bayes Factor analyses using a MATLAB package (Krekelberg, 2021).

## Results

We recorded MEG data while participants viewed a series of colored stimuli and performed either a categorization task or a discrimination task. During both tasks, reconstructed sources were focussed in visual cortex, and showed similar average event-related fields over time (**Figure 3**). To quantify the stimulus-related information present in the pattern of response across cortex, and to explore how neural representations of stimulus hue and luminance unfolded over time, we used a series of classification analyses.

**Figure 3:**
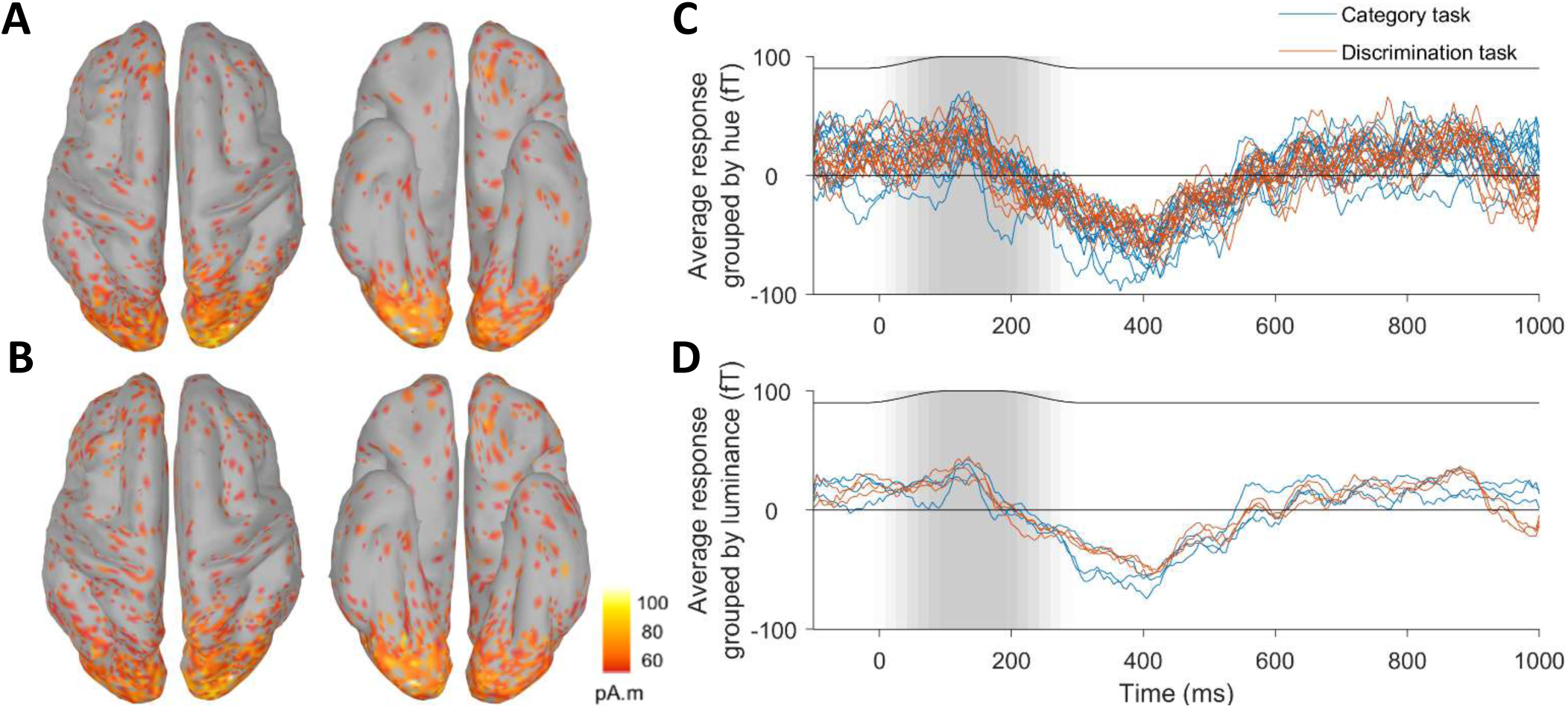
Estimated distributed source amplitudes (RMS values), averaged (n=8) for category (**A**) and discrimination (**B**) tasks, projected onto a partly inflated template cortical surface, viewed from above (left) and below (right). In **C** and **D**, the average response (event related fields) across 8 participants and all sensors is plotted, grouped by the 14 stimulus hues (**C**) or 3 luminance levels **(D**). Thin black lines in the upper part of each plot in **C** and **D**, along with the shaded grey background, indicate the temporal envelope of the stimulus, relative to trial onset

### Classification of stimulus hue and luminance

In **Figure 4** we show the average classifier accuracy, collapsed across all pairwise comparisons, when decoding hue and luminance. Across the two tasks, we found qualitatively similar timecourses of average classifier decoding for both stimulus features, with a trend toward higher classifier accuracy in the category task.

**Figure 4:**
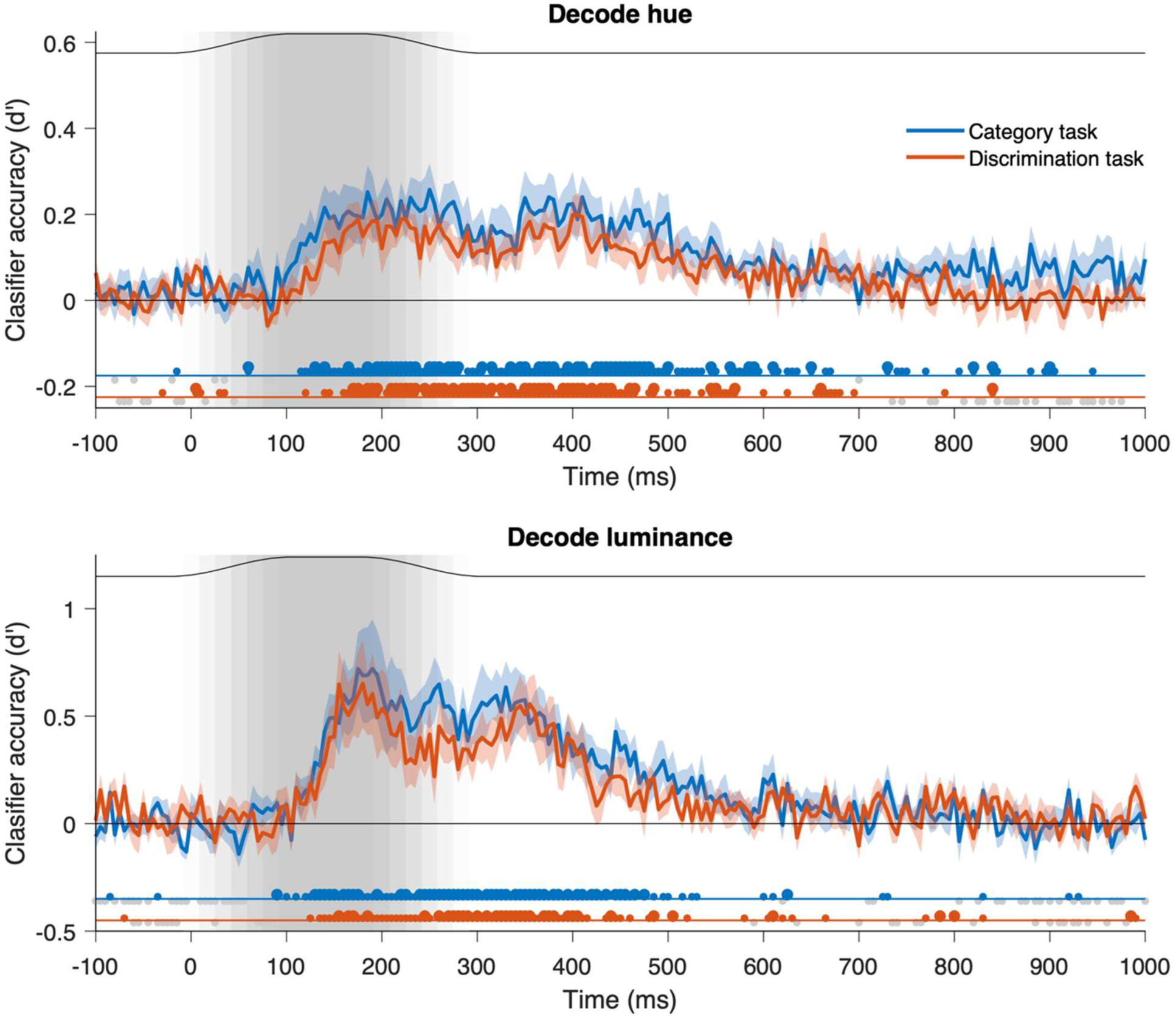
Decoding of stimulus hue (upper) and luminance (lower). Average classifier accuracy (n=8) across the two tasks is shown for classification of hue and luminance. Error bars are standard errors of the between-subject means. Thin black lines in the upper part of each plot, along with the shaded grey background, indicate the temporal envelope of the stimulus, relative to trial onset. Dots below the x-axis indicate the results of the Bayes Factor analyses. Colored dots show times when the classifier performance was above chance (small dots indicate moderate evidence BF>3, large dots indicate strong evidence, BF>10). Small grey dots indicate times where there was at least moderate evidence in favor of chance performance (BF<0.3).

This tendency for higher classifier accuracy in the category task may reflect the longer inter-stimulus interval for the category task (700ms) compared with the discrimination task (450ms), which we included to better match the tasks for difficulty (see Methods for details). The onset of above-chance decoding was approximately 100-120ms after stimulus onset in each case, which is slightly later than reported previously (e.g. Hermann et al., 2022). This later onset is likely because our stimuli did not onset abruptly but ramped on gradually, as shown in Figure 4.

#### Trend towards achromatic contrast, rather than intensity, in luminance decoding

Our stimuli were presented at three luminance levels: luminance decrements, stimuli isoluminant with the background, and luminance increments. If the neural responses were primarily driven by the luminance intensity differences between these, then discriminating increments vs decrements should lead to the highest classifier performance. However, if the neural responses to these stimuli were driven by luminance contrast more than intensity, then classifier performance should be highest when responses to the isoluminant stimuli are compared with either increments or decrements. These predictions are shown in the model DSMs (**Figure 5**, insets). Representational Similarity Analyses of the luminance decoding data are shown in **Figure 5**.

**Figure 5:**
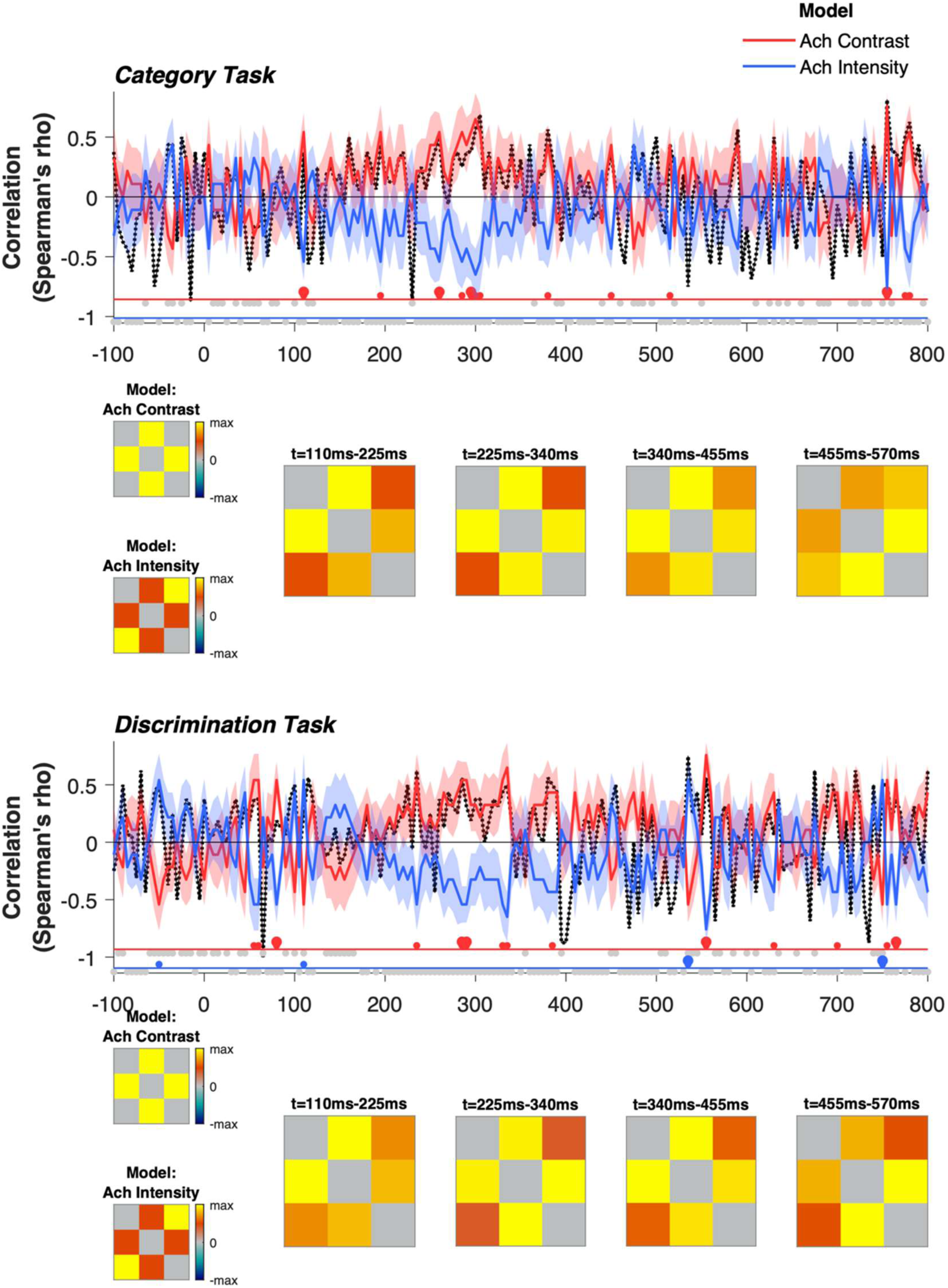
Representational Similarity Analysis of stimulus luminance, for the category (upper) and discrimination (lower) tasks. Average correlations (n=8) with the achromatic contrast and intensity models, along with the average dissimilarity matrices (DSMs) for four time windows (using the same time windows as in the reporting of hue RSA, below). Note that the achromatic contrast versus intensity models (shown in insets at the left) are perfectly negatively correlated, so the model correlations are mirror symmetrical about the x-axis (time axis) and positive correlation with one model will drive negative correlation with the other. The lower bound of the noise ceiling (Nili et al., 2014; see methods) is indicated by the dashed black line. Error bars are standard errors of the between-subject means, and dots below the x-axis indicate the results of the Bayes factor analysis, with plotting conventions as in Figure 4.

For both the category and the discrimination task, the data tended to be more consistent with the model based on achromatic contrast, rather than intensity. However, these effects were not particularly robust, as seen in the results of the Bayes Factor analysis in Figure 5. The average DSMs in Figure 5 are shown for four time windows to match those used for the hue analysis (see next section). Across these time windows, the average DSMs are consistent with the contrast model rather the intensity model across the first three time windows shown (110-455ms), with the lowest classifier performance for increments vs decrements in each case, but there was moderate variability across participants, which is seen in the standard error of the model correlations. The inter-participant variability is also seen in the noise ceiling, where the lower bound is not consistently above zero at time bins where there was above-chance classifier performance (as seen by comparing the dotted lines in Figure 5 with the average decoding of luminance in Figure 4). In these ways, these data are consistent with neural responses being driven by achromatic contrast, rather than intensity, but they provide only modest support for an effect in this direction.

### Dynamics of hue representations are consistent across task

Since each participant viewed stimuli of 14 different hues, the pairwise classifications of stimulus hue included 84 different comparisons. We include the average DSMs based on hue decoding for four time windows (see **Figure 6**). To better visualise the stimulus representations suggested by these DSMs, we also include MDS solutions based on each of these DSMs. When the DSMs are reduced with MDS, the resultant 2D solutions do not capture all information that may be present in the DSM. However, there is a reasonable *a priori* prediction that the neural representation of hue will be close to 2D, and the 14 stimulus hues can be represented in a number of different 2D spaces (such as the a*b* plane of L*a*b* space). In this way, the MDS solutions provide a useful insight into how the neural representations are structured if we assume that they are 2D. To test how well each MDS solution captured the corresponding DSM, in each case we rank-correlated (Spearman’s rho) the original DSM with a distance matrix defined by the best-fitting 2D MDS solution (Mantel test). These values were similarly high across time windows (see Figure 6).

**Figure 6:**
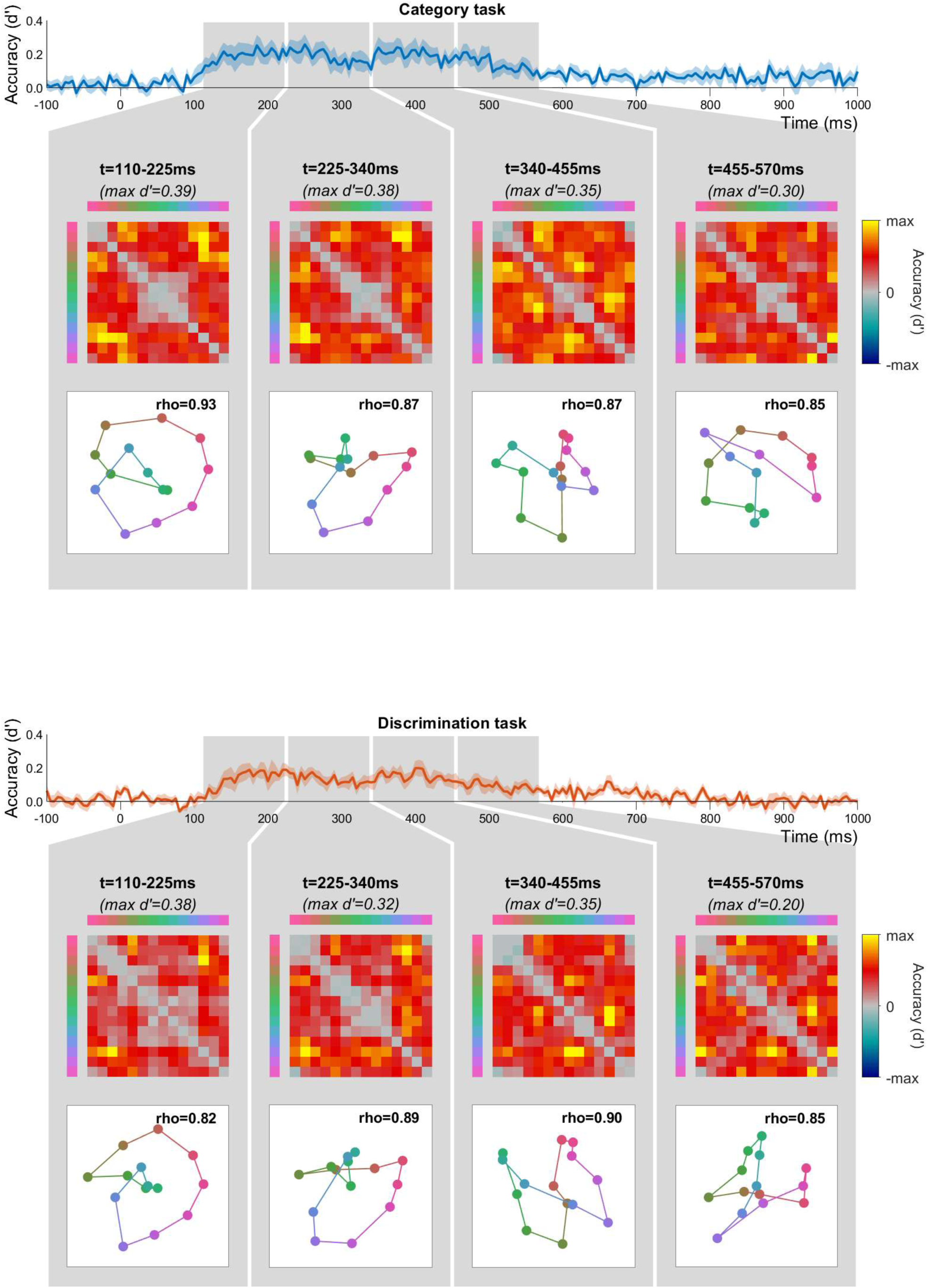
Average dissimilarity matrices (DSMs) and corresponding MDS solutions for data from the category task (upper) and discrimination task (lower). In each case, we considered data from four time windows of 115ms each, spanning the times that included above-chance classification of hue for both tasks. In both cases, the average decoding of hue (d’) is replotted from Figure 4, and grey shaded regions indicate the time windows corresponding to the four DSMs. The colormap used to plot each DSM is scaled by the maximum d’ of each matrix, indicated above each DSM. In the upper right of each MDS plot, rho values indicate the DSM/MDS correlation (Mantel test, see text for details).

We selected the four time windows in Figure 6 (also used in Figure 5) as equal-duration periods that together span the times with above-chance classification of stimulus hue. We chose to divide this period into four time windows in order to visualise the dynamics indicated by the RSA results (reported below), particularly the suggestion of a reorganisation at around 300-350ms after stimulus onset. For an alternate visualization, with shorter time windows that span the entire timecourse, we also include movies of the MDS solutions for each 50ms time window in the Appendix.

In comparing the DSMs and MDS solutions in Figure 6 with the model predictions in Figure 2, none of the time windows in either task provide a clear match to any of the 3 *a priori* models. Nonetheless, there are several features of the data that suggest that they reflect meaningful structure and can provide insight into how the neural representation of hue unfolds over time, including features that concur with one or more of the *a priori* models. For instance, all three models predict that classifier performance will tend to increase as hue difference increases, and this is consistent with all observed DSMs, which have relatively low classifier performance (close to grey) around their negative diagonals. In the MDS solutions, this is seen in tendency for hues to be arranged in order according to physical similarity (albeit with some clustering and/or folding).

For both the category task and discrimination task data, there is an evolution of the hue representation over time, as seen in the differences across the 4 time windows in Figure 6. The most striking feature of the data in Figure 6 is that this evolution across time windows is remarkably consistent for data from different tasks, particularly across the first 3 time windows. This is notable given that data for the two tasks were collected across different MEG recordings. This replication indicates a reliable but dynamic structure in the neural representations of hue, reflecting a reorganization of hue responses over time.

In the first two time windows of Figure 6, there is relatively low classifier performance amongst green hues: this is seen in the squares of grey in the DSMs, and the clustering of the greens in the MDS solutions. Interestingly, the greens do not remain clustered across the entire stimulus-induced response but show greater spacing in the third time window of Figure 6. This pattern is replicated across both tasks. In the third time window, MDS solutions from both tasks include a more substantial deviation from a circular arrangement, forming a ‘figure of 8’ with a crossing point around blue/yellow. Although the exploratory nature of these analyses precludes strong conclusions, they present interesting avenues for future work, as discussed in the general discussion below.

### RSA suggests a ‘coarse to fine’ transition in color representations

To better characterise how these hue representations change over the course of the stimulus-induced response, we performed a Representational Similarity Analysis, comparing the data with models based on cone contrast differences, hue difference, and color category (**Figure 7**) for data from both the discrimination and category tasks. All three models performed well overall (Figure 7A), with similar timing, which likely reflects the substantial overlap in their predictions based on their inter-model correlations (see Figure 2). Bayes Factors revealed moderate or strong effects across most times when classification of hue was above chance, and average model correlations that approached the lower-bound of the noise ceiling in each case. The strong performance of all three models we considered (Figure 7A) suggest a possible ceiling effect, where the shared predictions of these correlated models may account for performance approaching the lower-bound of the noise ceiling across the stimulus-induced response. These correlations offer limited additional insight into the dynamics of color evident in the dissimilarity matrices and MDS solutions (Figure 6).

**Figure 7:**
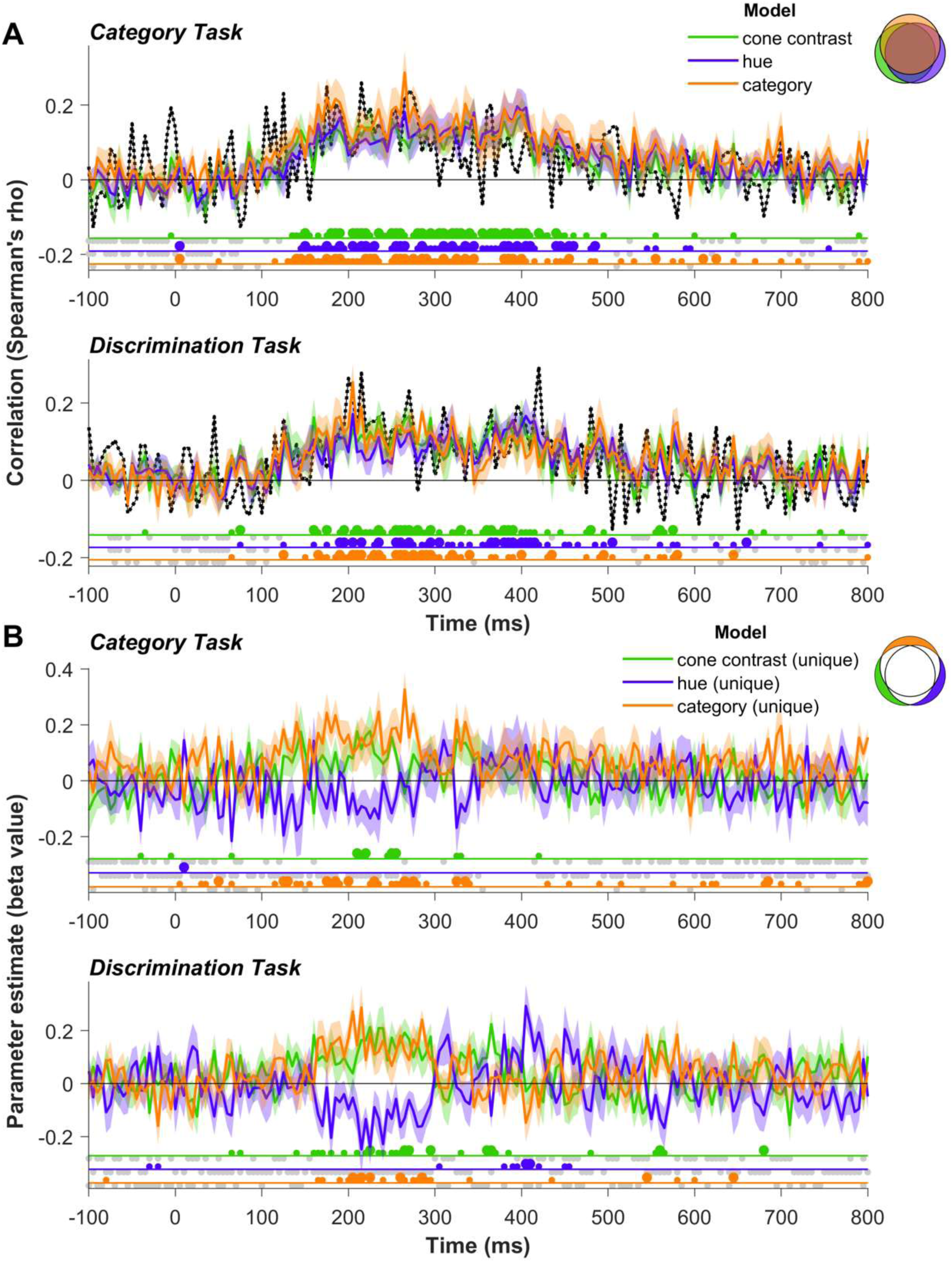
Representational similarity analysis. **A**: Average correlations between individual participants’ dissimilarity matrices for hue, and the cone contrast, hue and category models (see Figure 2 for models). **B**: Parameter estimates from GLM, quantifying the unique variance captured by each model, beyond the variance shared by other models (see methods for details). Colored regions of the Venn diagrams in the top right of **A** and **B** illustrate the variances captured by these different measures. In **A**, the lower bound of the noise ceiling (Nili et al., 2014; see methods) is indicated by the dashed black line. Error bars are standard errors of the between-subject means, and dots below the x-axis indicate the results of the Bayes factor analysis, with plotting conventions as in Figure 4.

To better isolate the predictions of each model from the others and test whether the unique variance captured by each model varies over time, we used a GLM analysis (detailed in the methods), with results shown in Figure 7**B**. The cone contrast model, which we expected to perform best for the earliest cortical responses, accounted for some unique variance in the first half of the response, but this was not clearly restricted to the response onset for either task. Interestingly, although we initially included the ‘category’ model to potentially capture later, higher-order color representations associated with color naming/categorization, we instead found that the unique variance captured by the category was most evident in the earlier part of the stimulus-induced response. This complements observations from the exploratory analysis (Figure 6) that the early colour responses included clustering of some hues of the same category, most notably the greens. For data from the discrimination task, we also found that the hue model captured variance beyond that of the category model later in the response. This suggests that later in the response, representations of similar hues in the same colour category are more distinct, reflecting their different appearance more than their category. This indication of a transition from the category to hue model better accounting for unique variance occurs at 300-350ms after stimulus onset, which is around the boundary between the second and third time windows in Figure 6. Together, these results suggest that the dynamic reorganization of colour representations could include a transition from a coarse, early response, with greater clustering of hues of the same category, to a representation that accentuates the finer appearance differences between within-category hues.

Despite very similar overall patterns of effects, the GLM analyses (Figure 7B) also suggest some task dependent effects, with the hue model accounting for unique variance at ∼400ms after stimulus onset when participants were performing the discrimination task, rather than the category task. The direction of this effect is consistent with the task requirements, since the discrimination task, but not the category task, required participants to pay attention to finer differences in hue, including within a colour category.

## Discussion

In this paper, we performed a series of classification analyses on the MEG recordings to explore how the neural response to colour unfolds over time, using a combination of exploratory analyses (MDS), and hypothesis-driven RSA. We chose stimuli with a relatively fine sampling of the hue space (14 hue directions) with the aim of revealing distortions of the neural representation of colour that could distinguish between different models of colour representation.

### The importance of comparing multiple models of colour representation

In the RSA, we tested three different representations (models) of stimulus colour, against the human cortical MEG data, with the aim of asking which model, if any, best captures the MEG-generated neural code. All models are based on human behavioural data. One representation of our chromatic stimuli is in a CIE L*a*b* space, where the fourteen suprathreshold colours are equally spaced around a colour circle in terms of their perceived colour differences (‘hue model’). In a second representation, these same stimuli are placed in a space based on the relative responses of the three different cone types to the stimuli (‘cone contrast model’). In this space, stimuli vary in a non-uniform way although retaining the overall colour stimulus ordering. The distortions in the cone contrast space reflect, at least in part, the differential contrast sensitivities of the two underlying cone opponent mechanisms, with the S-cone opponent direction elongated due to its poor contrast sensitivity compared to the highly sensitive L/M opponent mechanism (Cole et al., 1993; Sankeralli & Mullen, 1996). We selected a cone contrast model as it represents lower-level cortical responses in the visual system and provides a useful benchmark against which to compare other models. The simplest account of cortical colour representations would be that they reflect their subcortical inputs, so a robust test of higher-level models is whether they account for colour representations better than this lower-level one. The third colour representation we tested is based on the categorical groupings of our 14 colours, measured individually for each participant’s naming data, which produces a re-arrangement of the ‘hue model’ stimuli into clusters, predicting high within-category similarity as well as an orderly progression of categories based on their hue similarity (‘category model’).

### The challenge of overlapping models

These three models are conceptually distinct, and were selected as biologically plausible color spaces that may best capture the neural code. However, it is important to note that they are also correlated and so make substantially overlapping predictions in accounting for the observed data (e.g. the same ordering of hues). We overcome this issue by using the GLM analyses to compare the models in terms of the unique variance they capture (Figure 7B) which revealed differences between the three models to be seen that were invisible previously (Figure 7A). As mentioned above, comparing higher-level models with one based on a lower level of processing provides a valuable benchmark when considering what constitutes evidence in favor of a higher-level model, and for both hue and category models we find evidence that they significantly outperform the cone contrast model at different times.

### Robust decoding of luminance and hue

Across the stimulus-induced response, we found robust decoding of luminance and hue for both tasks, consistent with previous evidence that stimulus color can be decoded based on a topographic cortical representation of color in the MEG data (e.g. Goddard, Carlson, et al., 2022; Goddard, Shooner, et al., 2022; Hermann et al., 2022; Rosenthal et al., 2021; Teichmann et al., 2019, 2020). Decoding of both luminance and hue reached significance from approximately 100ms after initial stimulus onset and is optimal between 200 to 400ms. These onsets are slightly later than previously reported, and, unlike previous studies, we did not find evidence for decoding of luminance preceding that of hue (Hermann et al., 2022; Rosenthal et al., 2021). However, we note that our gradual onset stimuli may account for the later onset of the responses and are not optimised for comparing the onset of hue and luminance responses.

Patterns of luminance decoding (Figure 5) tended to indicate a neural representation based on luminance contrast, rather than luminance intensity. That is, accuracy was generally higher when the classifier was trained to discriminate between the isoluminant stimuli and those with luminance contrast, rather than between luminance increments and decrements, even though the latter have a greater difference in intensity. However, while decoding of luminance was highly robust, the pattern of decoding across luminance levels was variable across participants, and hence yielding only modest support for a neural representation of based on luminance contrast rather than intensity.

### Where in cortex are these hue and luminance representations located?

The spatial resolution of MEG source reconstruction does not allow us to make strong inferences about which visual cortical areas are contributing to the color representations. Although our source reconstruction indicated that sources were primarily located in visual cortex (Figure 3A-B), we found similar patterns of classifier results across early, ventral, and dorsal visual cortex (see Appendix). This could reflect dynamics that are propagating across these interconnected areas and/or that signals from these nearby regions were not well separated in our analyses. For this reason, the timing of the observed effects provides the clearest constraints on where these neural representations are located, which we consider below.

### Orderly hue representations, with minimal differences across task

RSA revealed hue representations that were less variable across participants than the luminance ones, possibly because our experimental design included larger variation in stimulus hue than luminance. From the onset of above-chance decoding, our data were consistent with all three of the models tested. This suggests that, from the earliest cortical responses to color, there is an orderly representation of hue. If the earliest stimulus-induced responses found here reflect the feedforward information in lower visual areas, then our data differ from fMRI studies showing that the responses of areas such as V1 and V2 lack an orderly representation of hue, with orderly responses emerging only in higher areas such as hV4 and VO (Brouwer & Heeger, 2009). This discrepancy could be related to the fact that, in the fMRI study, participants’ attention was diverted from the colored stimuli. Alternatively, it could be that any orderly representation of hue in V1 was relatively transient in the fMRI study, and beyond the temporal resolution of fMRI. The data presented here are consistent with more recent evidence from optical imaging suggesting a ‘pinwheel’ spatial organization of hue responses in V1 (Li et al., 2022).

The hue representations found here showed interesting temporal dynamics, considered below, but only modest variation with task. Using fMRI, Brouwer and Heeger (2013) demonstrated that areas hV4 and VO1 showed evidence of categorical clustering in hue representations, but this was present only when participants were engaged in a color naming task, and not when their attention was diverted. We were interested in whether the categorical clustering reported by Brouwer and Heeger (2013) was specific to a task requiring categorization, or a more general effect of attending to stimulus color. For this reason, we included a task requiring attention to color category (color-name 1-back task) as well as a task that required attention to color but without requiring categorization (exact-match 1-back task). The MDS and RSA analyses suggested that hue representations were largely consistent across the two tasks, but there was one noticeable trend in the RSA results in the direction of a task-specific effect, which we consider in the following section.

### Dynamics of hue representations, and a ‘coarse to fine’ transition

Our results show some interesting dynamics in the cortical response to color that are robust and replicated across data from both tasks. These dynamics are evident both in the MDS solutions (see Figure 6 or movies in the Appendix), and in the GLM analyses (Figure 7B). The GLM analyses, key to comparing the different models, indicate that there is a reorganization of the representation of color at around 300-350ms after stimulus-onset. Prior to this time, the data tend to be more consistent with the cone contrast model, potentially reflecting responses that modulate with cone contrast, and also with unique variance captured by the category model, supporting a representation based on a clustering of within-category hues. In the group data, this is seen in the MDS solutions primarily as a clustering of the neural responses to the green hues, although the dominance of the greens here may be driven by the fact that in our stimulus set, green was the largest category. After this reorganization, there is a more even separation of hues within the same category, as seen in the unique variance captured by the hue model in the latter part of the response for the discrimination task.

Inspection of the MDS solutions suggests that the neural data has systematic deviations from each of the three models we considered. Of course, it is possible that there are other color representations that would more closely match the observed data than the models we considered. Although we used the CIE L*a*b* space as an approximate model of color appearance in our ‘hue’ model, there are numerous models of color appearance based on apparent color difference or discriminability (Fairchild, 2005), or on natural image statistics (Smet et al., 2016) which would make similar predictions. However, given the substantial reorganization of these representations over time it is unlikely that there is any single (static) model that could capture the neural representational space across time. For instance, our ‘hue’ model may not be an accurate model of perceptual difference, since although CIE L*a*b* space is designed to be approximately perceptually uniform, it contains known deviations from a truly Euclidian space (Ennis & Zaidi, 2019). However, we find more pronounced deviations from the model based on CIE L*a*b* space than predicted by these known nonlinearities, and more importantly, the MDS results suggest that the neural data deviate from the CIE L*a*b* space in different ways over time: while greens are initially more clustered together than predicted by the hue model, at later times they are relatively more separated than other hues. Overall, the most pronounced feature of the hue representations in our data is their dynamic reorganization over time.

It is interesting that the category model performs best in the earlier part of the neural response. Since higher-level areas like hV4 and VO are associated with higher-level responses to color, including categorical representations (Brouwer & Heeger, 2013), we might expect that any categorical effects would emerge later in the stimulus-induced response, possibly incorporating feedback effects related to task and/or language. Instead, we found early contributions of the category model, meaning it is very unlikely that these reflect a higher-level representation where hues of the same semantic label are coded in a more similar way than hues of different names. Instead, we interpret these findings as suggesting that during earliest cortical responses there are hues which evoke very similar responses, which are then differentiated later in the response, but that this early clustering is not related to the influence of language. The ‘earlier’ timeframe of the categorial model includes responses from response onset, likely driven by areas such as V1 and V2, up to 300ms after stimulus onset, by which time the data would also include responses from higher-order areas such as hV4 and VO, and more anterior color centres (Lafer-Sousa et al., 2016; Zeki & Bartels, 1999). For instance, subdural electrode recordings in human ventral visual cortex, anterior to hV4, showed neuronal responses to color from ∼100ms after stimulus onset, peaking at ∼150ms after stimulus onset (Murphey et al., 2008).

After ∼300ms post stimulus-onset, there is no evidence of increasing categorical clustering. Hence, our data argue against the idea that any feedback or recurrent processing within cortex is driving hue representations to become more categorical over time. Instead, we find that hue representations of finer, within-category differences become stronger over time. This transition may echo the ‘coarse to fine’ strategy in different aspects of spatial vision, with the coarse color structure processed before the fine within-color detail. While coarse-to-fine processing typically refers to neural responses shifting from lower to higher spatial scale information over time (e.g. Goffaux et al., 2011; Watt, 1987), equivalent coding principles have been implicated in orientation coding (Ringach et al., 1997, 2003; Shapley et al., 2003), and face processing (Dobs et al., 2019), where initial responses are more coarsely tuned, preceding representations of finer differences (i.e. wider then narrower tuning for orientation responses, face age and gender preceding representations that differentiate between face identities).

The late time at which we observe this transition to an enhanced representation of finer hue differences (∼300ms) suggests that these effects could be at least partly driven by top-down effects of attention, which have previously been reported to produced effects consistent with sharpening of population tuning for color (Bartsch et al., 2017; Goddard, Carlson, et al., 2022). In both conditions, participants were attending to color, and across both tasks we saw similar dynamics, consistent with a transition from coarser to finer hue representations. However, at later stages of the stimulus induced response there was stronger evidence in favor of the hue model for data from the discrimination task than for the category task at corresponding times (Figure 7B). This task-specific effect is much subtler than the fMRI differences when comparing a categorical task with diverted attention (Brouwer & Heeger, 2013), suggesting that the effect of attending to a colored stimulus, rather than specifically attending to its category, is of greater relevance.

A final notable feature of the hue representations in our data is in the latter part of the neural response (e.g. the 340-455ms and 455-570ms time windows in Figure 6), the hue closest to yellow is also close to one or more blue hues. This is clearest in the MDS results in the 340-455ms time window, where the hue circle is represented more as a ‘figure of 8’, with crossing at yellow/blue, but within greenish and reddish hues there is greater separation. This is not accounted for by the S-cone inputs being slower to reach cortex than red-green signals carried by the parvocellular pathway (e.g. Cottaris & De Valois, 1998), since the effect we observe is well after the initial cortical response, and because the effect is in the opposite direction, with a relatively late encoding of red-green differences. Although these observations are still speculative and based on an exploratory analysis, we note that in both tasks, the yellow/blue crossing occurred not for the extremes along the S-cone isolating axis, but closer to unique yellow/blue, and the daylight axis (Delahunt & Brainard, 2004). We did not design this work to test for such asymmetries, but these observations could motivate future work to characterise these.

## Conclusion

In summary, in this MEG study we reveal a dynamic reorganization of the cortical response to color over time, which is remarkably consistent across our two tasks. We compared three conceptually distinct, biologically plausible models that may best capture the neural code, using GLM analyses to isolate the unique data variance accounted for by each model. The resulting model dynamics reflects a ‘coarse to fine’ transition over time from an early categorial grouping of colors in the cortical representation evolving into to a finer, more even separation of the hues.

## Acknowledgements

This work was funded by Canadian Institutes of Health Research (CIHR) grants 153277 and 10819 to KTM. EG was partly supported by the Australian Research Council (DP220100747). We thank Christopher Shooner for help with the design and data collection, and Marc Lalancette for advice on MEG protocols. We thank all our participants for their assistance with data collection.

## Data Availability

Data from MEG experiments are freely available online from the Open Science Framework (https://doi.org/10.17605/OSF.IO/KMFZ7). This online repository includes deidentified MEG data from the MEG experiments and details of the stimulus timing for each participant.

## Notes

### Competing Interest Statement

The authors have declared no competing interest.

### Summary of Updates

Cross-temporal decoding analysis removed, additional visualisation of univariate responses added.

## References

Bae, G.-Y., & Chen, K.-W. (2024). EEG decoding reveals task-dependent recoding of sensory information in working memory. NeuroImage, 297, 120710. 10.1016/j.neuroimage.2024.120710

Bartsch, M. V., Loewe, K., Merkel, C., Heinze, H.-J., Schoenfeld, M. A., Tsotsos, J. K., & Hopf, J.-M. (2017). Attention to Color Sharpens Neural Population Tuning via Feedback Processing in the Human Visual Cortex Hierarchy. Journal of Neuroscience, 37(43), 10346–10357. 10.1523/JNEUROSCI.0666-17.2017

Berlin, B., & Kay, P. (1969). Basic color terms: Their universality and evolution. University of California Press.

Brainard, D. H. (1997). The Psychophysics Toolbox. Spatial Vision, 10(4), 433–436.

Brewer, A. A., Liu, J., Wade, A. R., & Wandell, B. A. (2005). Visual field maps and stimulus selectivity in human ventral occipital cortex. Nature Neuroscience, 8(8), 1102–1109. 10.1038/nn1507

Brouwer, G. J., & Heeger, D. J. (2009). Decoding and reconstructing color from responses in human visual cortex. Journal of Neuroscience, 29(44), 13992–14003.

Brouwer, G. J., & Heeger, D. J. (2013). Categorical clustering of the neural representation of color. Journal of Neuroscience, 33(39), 15454–15465. 10.1523/JNEUROSCI.2472-13.2013

Chauhan, T., Jakovljev, I., Thompson, L. N., Wuerger, S. M., & Martinovic, J. (2023). Decoding of EEG signals reveals non-uniformities in the neural geometry of colour. NeuroImage, 268, 119884. 10.1016/j.neuroimage.2023.119884

Cole, G. R., Hine, T., & McIlhagga, W. (1993). Detection mechanisms in L-, M-, and S-cone contrast space. Journal of the Optical Society of America A-Optics Image Science and Vision, 10(1), 38–51.

Conway, B. R., Malik-Moraleda, S., & Gibson, E. (2023). Color appearance and the end of Hering’s Opponent-Colors Theory. Trends in Cognitive Sciences, 27(9), 791–804. 10.1016/j.tics.2023.06.003

Cottaris, N. P., & De Valois, R. L. (1998). Temporal dynamics of chromatic tuning in macaque primary visual cortex. Nature, 395(6705), 896–900. 10.1038/27666

Dale, A. M., Fischl, B., & Sereno, M. I. (1999). Cortical surface-based analysis. I: Segmentation and surface reconstruction. Neuroimage, 9(2), 179–194.

de Valois, R. L., & de Valois, K. K. (1993). A multi-stage color model. Vision Research, 33(8), 1053–1065.

Delahunt, P. B., & Brainard, D. H. (2004). Does human color constancy incorporate the statistical regularity of natural daylight? Journal of Vision, 4(2), 57–81.

Derrington, A. M., Krauskopf, J., & Lennie, P. (1984). Chromatic mechanisms in lateral geniculate nucleus of macaque. Journal of Physiology, 357, 241–265.

Dobs, K., Isik, L., Pantazis, D., & Kanwisher, N. (2019). How face perception unfolds over time. Nature Communications, 10(1), Article 1. 10.1038/s41467-019-09239-1

Engel, S. A., Rumelhart, D. E., Wandell, B. A., Lee, A. T., Glover, G. H., Chichilnisky, E. J., & Shadlen, M. N. (1994). fMRI of human visual cortex. Nature, 369(6481), 525.

Ennis, R. J., & Zaidi, Q. (2019). Geometrical structure of perceptual color space: Mental representations and adaptation invariance. Journal of Vision, 19(12), 1. 10.1167/19.12.1

Eskew, R. T. (2009). Higher-order color mechanisms: A critical review. Vision Research, 49(22), 2686–2704.

Fairchild, M. D. (2005). Color Appearance Models. John Wiley & Sons, Inc.

Farnsworth, D. (1957). The Farnsworth-Munsell 100-hue test for the examination of color discrimination. Macbeth, a division of Kollmorgen Corp.

Fischl, B., Sereno, M. I., & Dale, A. M. (1999). Cortical surface-based analysis. II: Inflation, flattening, and a surface-based coordinate system. Neuroimage, 9(2), 195–207.

Goddard, E., Carlson, T. A., & Woolgar, A. (2022). Spatial and feature-selective attention have distinct, interacting, effects on population-level tuning. J Cogn Neurosci, 34(2), 290–312. 10.1162/jocn_a_01796

Goddard, E., Chang, D. H. F., Hess, R. F., & Mullen, K. T. (2019). Color contrast adaptation: fMRI fails to predict behavioral adaptation. NeuroImage, 201, 116032. 10.1016/j.neuroimage.2019.116032

Goddard, E., Mannion, D. J., McDonald, J. S., Solomon, S. G., & Clifford, C. W. G. (2011). Color responsiveness argues against a dorsal component of human V4. Journal of Vision, 11(4), 3, 1–21. 10.1167/11.4.3

Goddard, E., Mannion, D. M., McDonald, J. S., Solomon, S. G., & Clifford, C. W. G. (2010). Combination of Subcortical Color Channels in Human Visual Cortex. Journal of Vision, 10(5), 25, 1–17. 10.1167/10.5.25

Goddard, E., & Mullen, K. T. (2020). fMRI representational similarity analysis reveals graded preferences for chromatic and achromatic stimulus contrast across human visual cortex. NeuroImage, 215, 116780. 10.1016/j.neuroimage.2020.116780

Goddard, E., & Mullen, K. T. (2021). Attention selectively enhances stimulus information for surround over foveal stimulus representations in occipital cortex. Journal of Vision, 21(3), 20. 10.1167/jov.21.3.20

Goddard, E., Shooner, C., & Mullen, K. T. (2022). Magnetoencephalography contrast adaptation reflects perceptual adaptation. Journal of Vision, 22(10), 16, 1–19. 10.1167/jov.22.10.16

Goffaux, V., Peters, J., Haubrechts, J., Schiltz, C., Jansma, B., & Goebel, R. (2011). From coarse to fine? Spatial and temporal dynamics of cortical face processing. Cerebral Cortex, 21(2), 467–476. 10.1093/cercor/bhq112

Golub, G. H., & Loan, C. F. V. (1996). Matrix Computations, Third Edition. Johns Hopkins University Press.

Grootswagers, T., Robinson, A. K., Shatek, S. M., & Carlson, T. A. (2024). Mapping the dynamics of visual feature coding: Insights into perception and integration. PLOS Computational Biology, 20(1), e1011760. 10.1371/journal.pcbi.1011760

Hajonides, J. E., Nobre, A. C., van Ede, F., & Stokes, M. G. (2021). Decoding visual colour from scalp electroencephalography measurements. NeuroImage, 118030. 10.1016/j.neuroimage.2021.118030

He, J., Taveras Cruz, Y., & Eskew, R. T., Jr. (2020). Methods for determining equiluminance in terms of L/M cone ratios. Journal of Vision, 20(4), 22. 10.1167/jov.20.4.22

Hermann, K. L., Singh, S. R., Rosenthal, I. A., Pantazis, D., & Conway, B. R. (2022). Temporal dynamics of the neural representation of hue and luminance polarity. Nature Communications, 13(1), 661. 10.1038/s41467-022-28249-0

Huang, M. X., Mosher, J. C., & Leahy, R. M. (1999). A sensor-weighted overlapping-sphere head model and exhaustive head model comparison for MEG. Physics in Medicine and Biology, 44, 423–440. 10.1088/0031-9155/44/2/010

Huk, A., Dougherty, R., & Heeger, D. (2002). Retinotopy and functional subdivision of human areas MT and MST. Journal of Neuroscience, 22(16), 7195–7205.

Ishihara, S. (1990). Ishihara’s tests for color-blindness, 38 plate ed. Kanehara, Shuppan Co. Ltd.

Kass, R. E., & Raftery, A. E. (1995). Bayes Factors. Journal of the American Statistical Association, 90(430), 773–795. 10.1080/01621459.1995.10476572

Kleiner, M., Brainard, D., & Pelli, D. G. (2007). What’s new in Psychtoolbox-3? Perception, 36, ECVP Abstract Supplement.

Komatsu, H., Ideura, Y., Kaji, S., & Yamane, S. (1992). Color selectivity of neurons in the inferior temporal cortex of the awake macaque monkey. Journal of Neuroscience, 12(2), 408–424. 10.1523/JNEUROSCI.12-02-00408.1992

Krekelberg, B. (2021). *klabhub/bayesFactor: Ttest updates* (Version v2.2.0) [Computer software]. Zenodo. 10.5281/zenodo.5707551

Kriegeskorte, N., Mur, M., & Bandettini, P. (2008). Representational similarity analysis—Connecting the branches of systems neuroscience. Frontiers in Systems Neuroscience, 2, 4. 10.3389/neuro.06.004.2008

Kriegeskorte, N., Mur, M., Ruff, D. A., Kiani, R., Bodurka, J., Esteky, H., Tanaka, K., & Bandettini, P. A. (2008). Matching categorical object representations in inferior temporal cortex of man and monkey. Neuron, 60(6), 1126–1141. 10.1016/j.neuron.2008.10.043

Lafer-Sousa, R., & Conway, B. R. (2013). Parallel, multi-stage processing of colors, faces and shapes in macaque inferior temporal cortex. Nature Neuroscience, 16(12), 1870–1878. 10.1038/nn.3555

Lafer-Sousa, R., Conway, B. R., & Kanwisher, N. G. (2016). Color-Biased Regions of the Ventral Visual Pathway Lie between Face- and Place-Selective Regions in Humans, as in Macaques. Journal of Neuroscience, 36(5), 1682–1697. 10.1523/JNEUROSCI.3164-15.2016

Larsson, J., & Heeger, D. J. (2006). Two retinotopic visual areas in human lateral occipital cortex. Journal of Neuroscience, 26(51), 13128–13142.

Lennie, P., & Movshon, J. A. (2005). Coding of color and form in the geniculostriate visual pathway. Journal of the Optical Society of America A-Optics Image Science and Vision, 22(10), 2013–2033.

Li, P., Garg, A. K., Zhang, L. A., Rashid, M. S., & Callaway, E. M. (2022). Cone opponent functional domains in primary visual cortex combine signals for color appearance mechanisms. Nature Communications, 13(1), 6344. 10.1038/s41467-022-34020-2

Michna, M. L., Yoshizawa, T., & Mullen, K. T. (2007). S-cone contributions to linear and non-linear motion processing. Vision Research, 47(8), 1042–1054. 10.1016/j.visres.2007.01.014

Morey, R. D., & Wagenmakers, E.-J. (2014). Simple relation between Bayesian order-restricted and point-null hypothesis tests. Statistics & Probability Letters, 92, 121–124. 10.1016/j.spl.2014.05.010

Mullen, K. T., Dumoulin, S. O., & Hess, R. F. (2008). Color responses of the human lateral geniculate nucleus: Selective amplification of S-cone signals between the lateral geniculate nucleno and primary visual cortex measured with high-field fMRI. European Journal of Neuroscience, 28(9), 1911–1923.

Mullen, K. T., Dumoulin, S. O., McMahon, K. L., de Zubicaray, G. I., & Hess, R. F. (2007). Selectivity of human retinotopic visual cortex to S-cone-opponent, L/M-cone-opponent and achromatic stimulation. European Journal of Neuroscience, 25(2), 491–502. 10.1111/j.1460-9568.2007.05302.x

Mullen, K. T., & Sankeralli, M. J. (1999). Evidence for the stochastic independence of the blue-yellow, red-green and luminance detection mechanisms revealed by subthreshold summation. Vision Research, 39, 733–745.

Murphey, D. K., Yoshor, D., & Beauchamp, M. S. (2008). Perception matches selectivity in the human anterior color center. Current Biology, 18(3), 216–220.

Nili, H., Wingfield, C., Walther, A., Su, L., Marslen-Wilson, W., & Kriegeskorte, N. (2014). A toolbox for representational similarity analysis. PLoS Computational Biology, 10(4), e1003553. 10.1371/journal.pcbi.1003553

Parkes, L. M., Marsman, J. C., Oxley, D. C., Goulermas, J. Y., & Wuerger, S. M. (2009). Multivoxel fMRI analysis of color tuning in human primary visual cortex. Journal of Vision, 9(1), 1, 1–13.

Pelli, D. G. (1997). The VideoToolbox software for visual psychophysics: Transforming numbers into movies. Spatial Vision, 10(4), 437–442.

Popal, H., Wang, Y., & Olson, I. R. (2019). A Guide to Representational Similarity Analysis for Social Neuroscience. Social Cognitive and Affective Neuroscience, 14(11), 1243–1253. 10.1093/scan/nsz099

Retter, T. L., Gao, Y., Jiang, F., Rossion, B., & Webster, M. A. (2023). Automatic, Early Color-Specific Neural Responses to Object Color Knowledge. Brain Topography, 36(5), 710–726. 10.1007/s10548-023-00979-4

Ringach, D. L., Hawken, M. J., & Shapley, R. (1997). Dynamics of orientation tuning in macaque primary visual cortex. Nature, 387(6630), 281–284.

Ringach, D. L., Hawken, M. J., & Shapley, R. (2003). Dynamics of Orientation Tuning in Macaque V1: The Role of Global and Tuned Suppression. Journal of Neurophysiology, 90(1), 342–352. 10.1152/jn.01018.2002

Rosenthal, I. A., Singh, S. R., Hermann, K. L., Pantazis, D., & Conway, B. R. (2021). Color Space Geometry Uncovered with Magnetoencephalography. Current Biology: CB, 31(3), 515–526.e5. 10.1016/j.cub.2020.10.062

Rozman, A., Chauhan, T., & Martinovic, J. (2024). Characterising Representations of Hue and Saturation in the Cortex Using Information Decoding (p. 2024.12.19.628842). bioRxiv. 10.1101/2024.12.19.628842

Sandhaeger, F., von Nicolai, C., Miller, E. K., & Siegel, M. (2019). Monkey EEG links neuronal color and motion information across species and scales. eLife, 8. 10.7554/eLife.45645

Sankeralli, M. J., & Mullen, K. T. (1996). Estimation of the L-, M-, and S-cone weights of the postreceptoral detection mechanisms. JOSA A, 13(5), 906–915.

Schira, M. M., Tyler, C. W., Breakspear, M., & Spehar, B. (2009). The foveal confluence in human visual cortex. Journal of Neuroscience, 29(28), 9050–9058.

Sereno, M. I., Dale, A. M., Reppas, J. B., Kwong, K. K., Belliveau, J. W., Brady, T. J., Rosen, B. R., & Tootell, R. B. (1995). Borders of multiple visual areas in humans revealed by functional magnetic resonance imaging. Science, 268(5212), 889–893.

Shapley, R., Hawken, M., & Ringach, D. L. (2003). Dynamics of orientation selectivity in the primary visual cortex and the importance of cortical inhibition. Neuron, 38(5), 689–699.

Siuda-Krzywicka, K., Boros, M., Bartolomeo, P., & Witzel, C. (2019). The biological bases of colour categorisation: From goldfish to the human brain. Cortex, 118, 82–106. 10.1016/j.cortex.2019.04.010

Smet, K. A. G., Webster, M. A., & Whitehead, L. A. (2016). A simple principled approach for modeling and understanding uniform color metrics. Journal of the Optical Society of America. A, Optics, Image Science, and Vision, 33(3), A319–331. 10.1364/JOSAA.33.00A319

Smith, V. C., & Pokorny, J. (1975). Spectral sensitivity of the foveal cone photopigments between 400 and 500 nm. Vision Research, 15(2), 161–171.

Stockman, A., & Brainard, D. H. (2010). Color Vision Mechansims. In M. Bass (Ed.), *Handbook* of Optics: Volume III - Vision and Vision Optics (3rd Edition). McGraw-Hill Education. https://www.accessengineeringlibrary.com/content/book/9780071498913/chapter/chapter11

Stoughton, C. M., & Conway, B. R. (2008). Neural basis for unique hues. Current Biology, 18(16), R698–9.

Sutterer, D. W., Coia, A. J., Sun, V., Shevell, S. K., & Awh, E. (2021). Decoding chromaticity and luminance from patterns of EEG activity. Psychophysiology, 58(4), e13779. 10.1111/psyp.13779

Tadel, F., Baillet, S., Mosher, J. C., Pantazis, D., & Leahy, R. M. (2011). Brainstorm: A user-friendly application for MEG/EEG analysis. Computational Intelligence and Neuroscience, 2011, 879716. 10.1155/2011/879716

Teichmann, L., Grootswagers, T., Carlson, T., & Rich, A. N. (2019). Seeing versus knowing: The temporal dynamics of real and implied colour processing in the human brain. NeuroImage, 200(15), 373–381. 10.1016/j.neuroimage.2019.06.062

Teichmann, L., Moerel, D., Baker, C., & Grootswagers, T. (2022). An Empirically Driven Guide on Using Bayes Factors for M/EEG Decoding. Aperture Neuro, 2, 1–10. 10.52294/ApertureNeuro.2022.2.MAOC6465

Teichmann, L., Quek, G. L., Robinson, A. K., Grootswagers, T., Carlson, T. A., & Rich, A. N. (2020). The Influence of Object-Color Knowledge on Emerging Object Representations in the Brain. The Journal of Neuroscience: The Official Journal of the Society for Neuroscience, 40(35), 6779–6789. 10.1523/JNEUROSCI.0158-20.2020

Uusitalo, M. A., & Ilmoniemi, R. J. (1997). Signal-space projection method for separating MEG or EEG into components. Medical & Biological Engineering & Computing, 35(2), 135–140. 10.1007/BF02534144

Watt, R. J. (1987). Scanning from coarse to fine spatial scales in the human visual system after the onset of a stimulus. JOSA A, 4(10), 2006–2021. 10.1364/JOSAA.4.002006

Witzel, C. (2019). Misconceptions About Colour Categories. Review of Philosophy and Psychology, 10(3), 499–540. 10.1007/s13164-018-0404-5

Wu, Y., Zhang, Y., Mao, Y., Feng, K., Wei, D., & Song, L. (2022). Reconstructing sources location of visual color cortex by the task-irrelevant visual stimuli through machine learning decoding. Heliyon, 8(12). 10.1016/j.heliyon.2022.e12287

Zeki, S., & Bartels, A. (1999). The clinical and functional measurement of cortical (in)activity in the visual brain, with special reference to the two subdivisions (V4 and V4 alpha) of the human colour centre. Philosophical Transactions of the Royal Society of London Series B-Biological Sciences, 354(1387), 1371–1382.

