## Appendix for "Temporal evolution of color representations measured with MEG reveals a ‘coarse to fine’ dynamic"

This appendix includes figures equivalent to those in the main text but for other regions of interest (ventral visual cortex, VVC, and dorsal visual cortex, DVC), in Figures A1-A8.

Directly below we include a description of the videos included in this appendix as separate files.

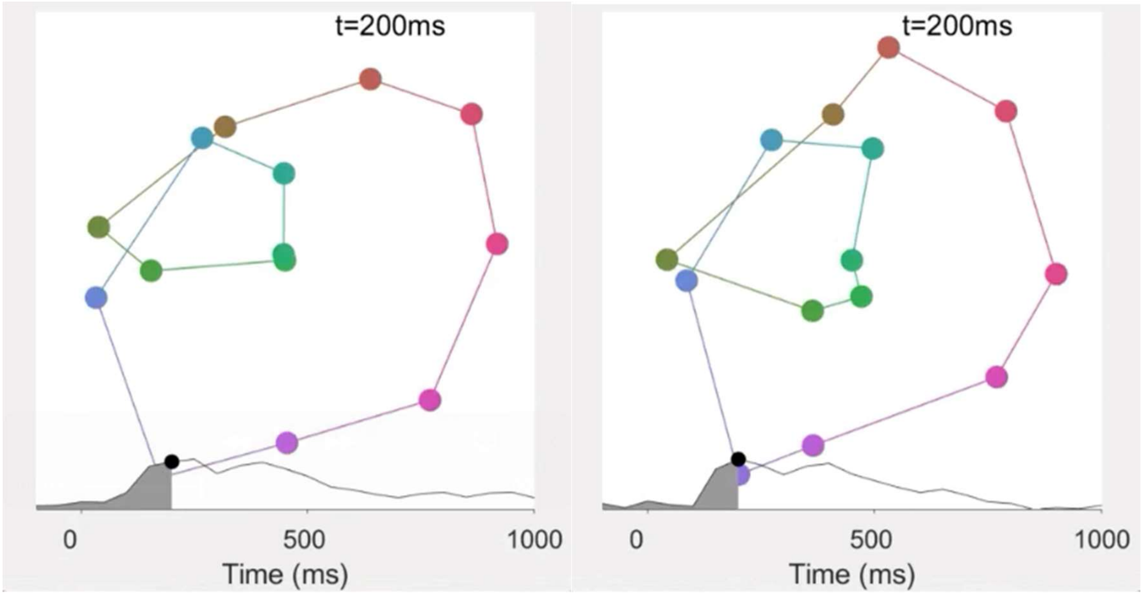


***Videos A1-A2****: Videos of MDS solutions for data from the category task (left) and discrimination task (right) over time (-100-1000ms, 50ms bins), based on classifier performance in the EVC (n=8). Note: above is a still frame from the videos, which are included in the appendix as separate files.*

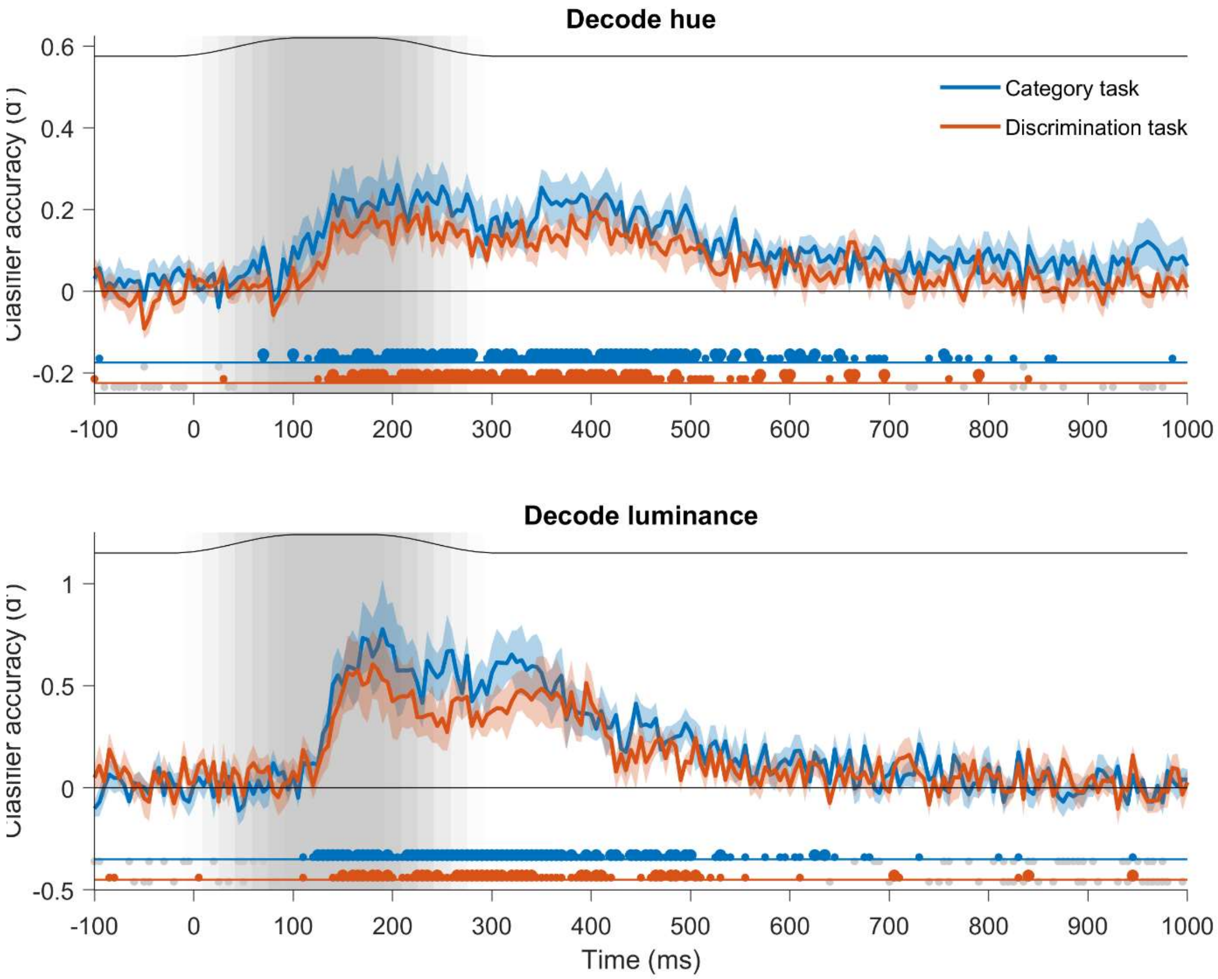


***Figure A1:*** *Decoding of stimulus hue (upper) and luminance (lower) for classifiers trained on data from the VVC. Plotting conventions as in Figure 4.*

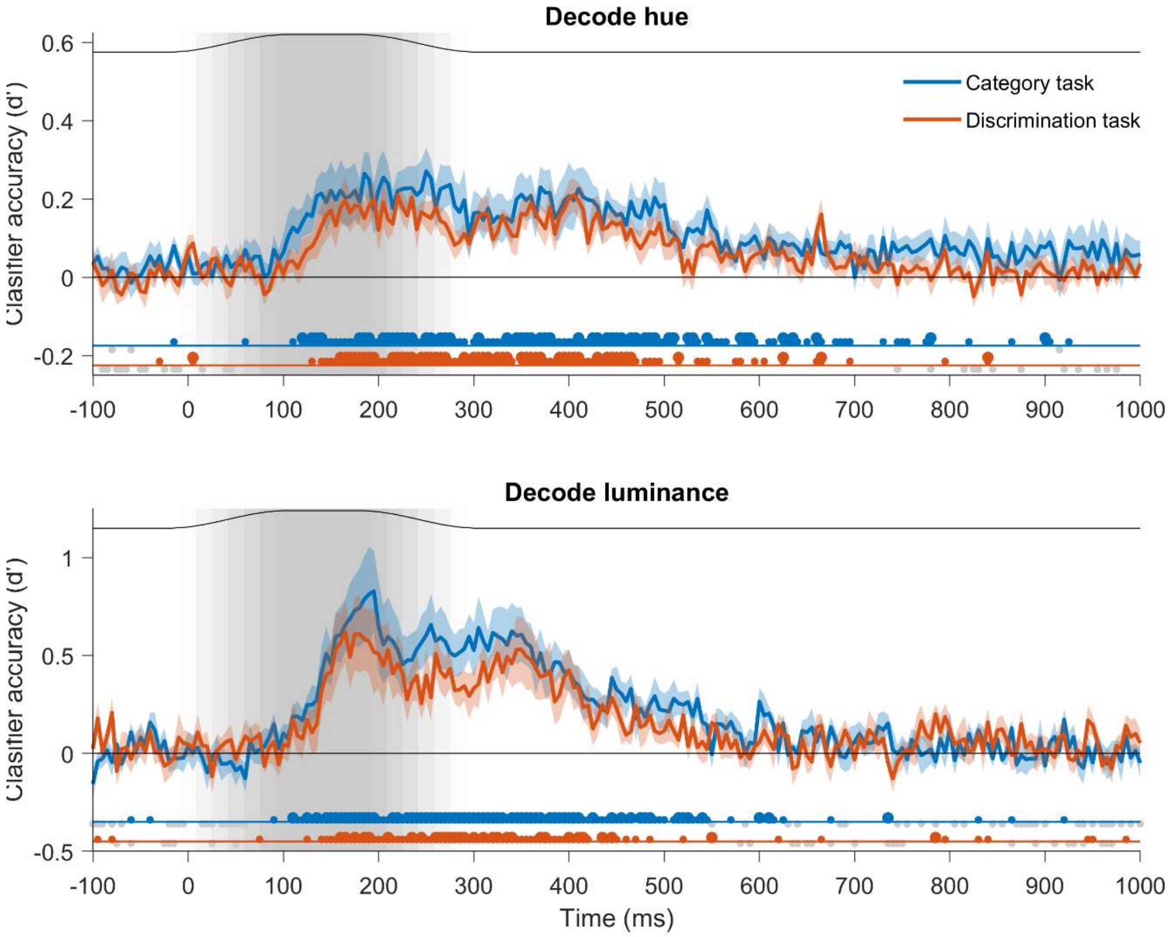


***Figure A2:*** *Decoding of stimulus hue (upper) and luminance (lower) for classifiers trained on data from the DVC. Plotting conventions as in Figure 4.*

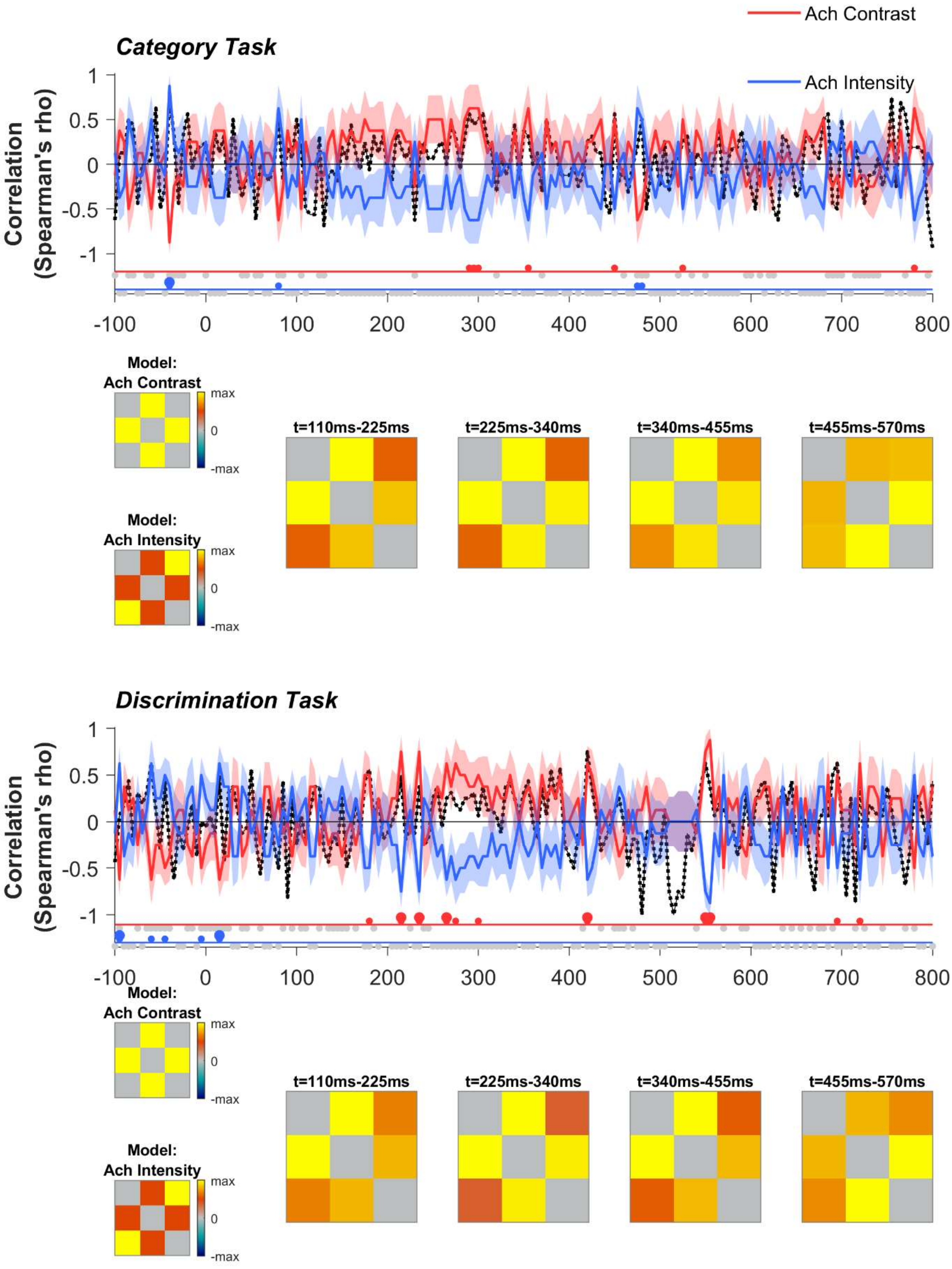


***Figure A3:*** *Representational Similarity Analysis of stimulus luminance, for the category (upper) and discrimination (lower) tasks for classifiers trained on data from the VVC. Plotting conventions as in Figure 5.*

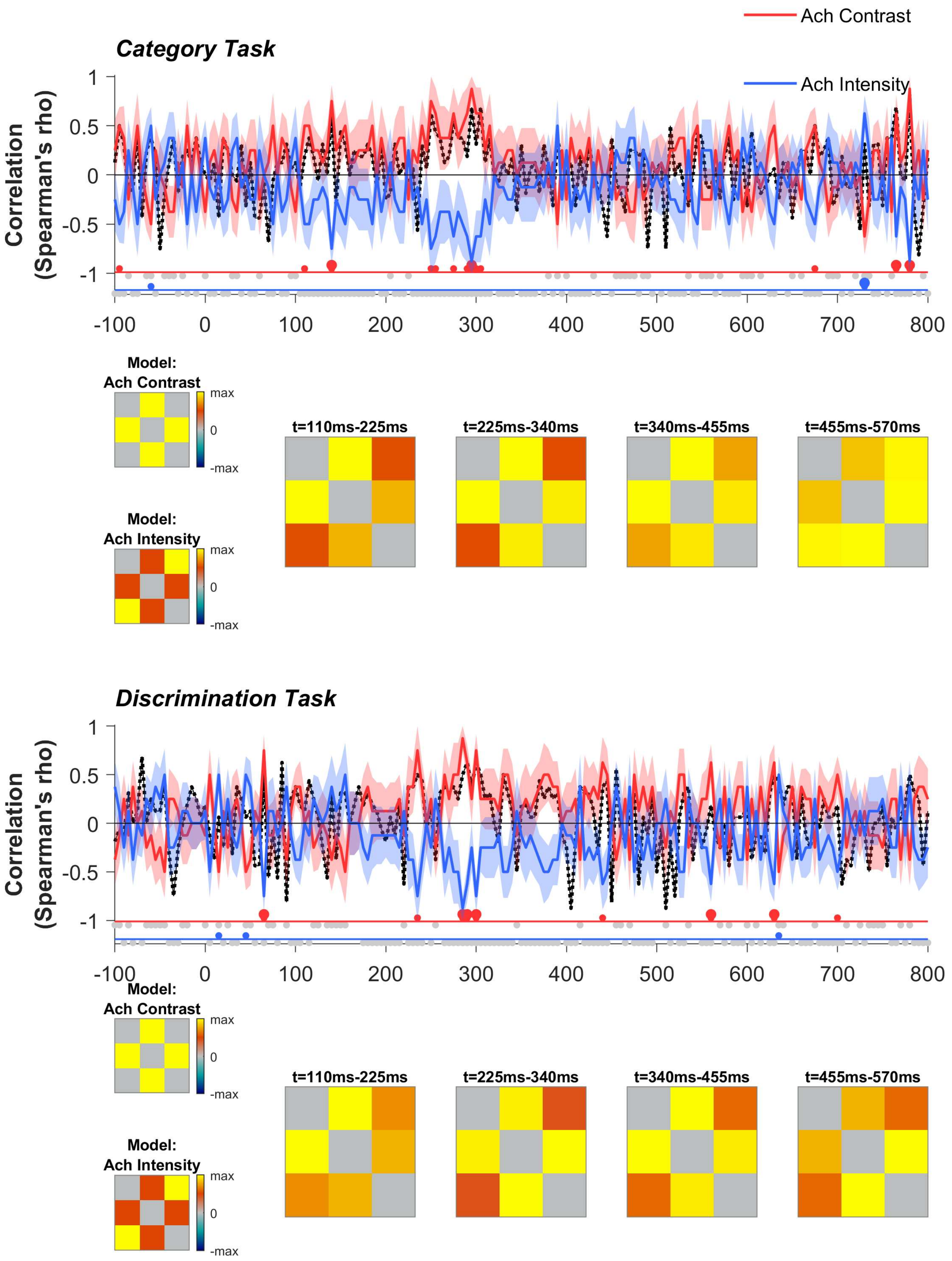


***Figure A4:*** *Representational Similarity Analysis of stimulus luminance, for the category (upper) and discrimination (lower) tasks for classifiers trained on data from the DVC. Plotting conventions as in Figure 5.*

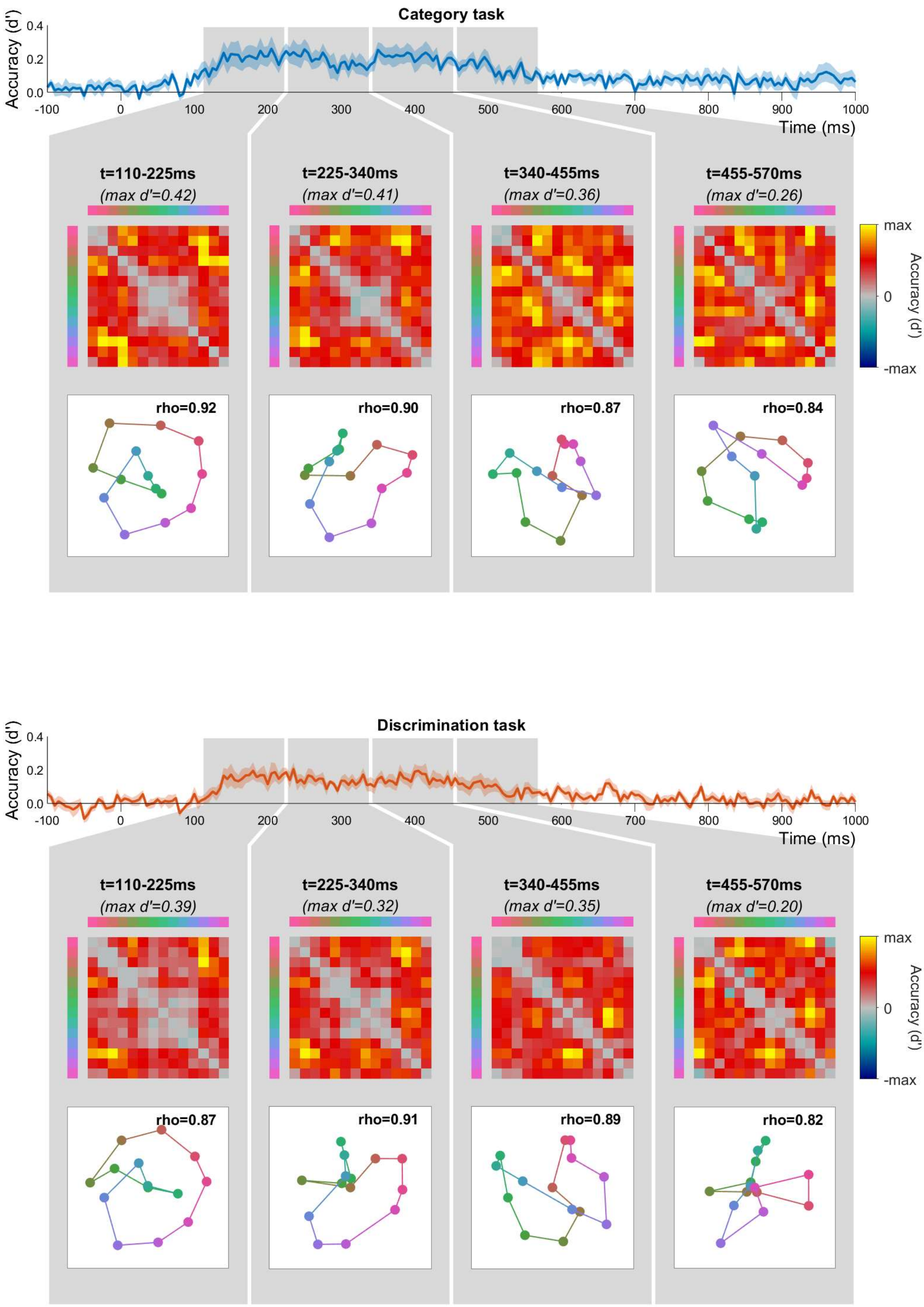


***Figure A5***: *Average dissimilarity matrices (DSMs) and corresponding MDS solutions for data from the category task (upper) and discrimination task (lower) for classifiers trained on data from the VVC. Plotting conventions as in Figure 6.*

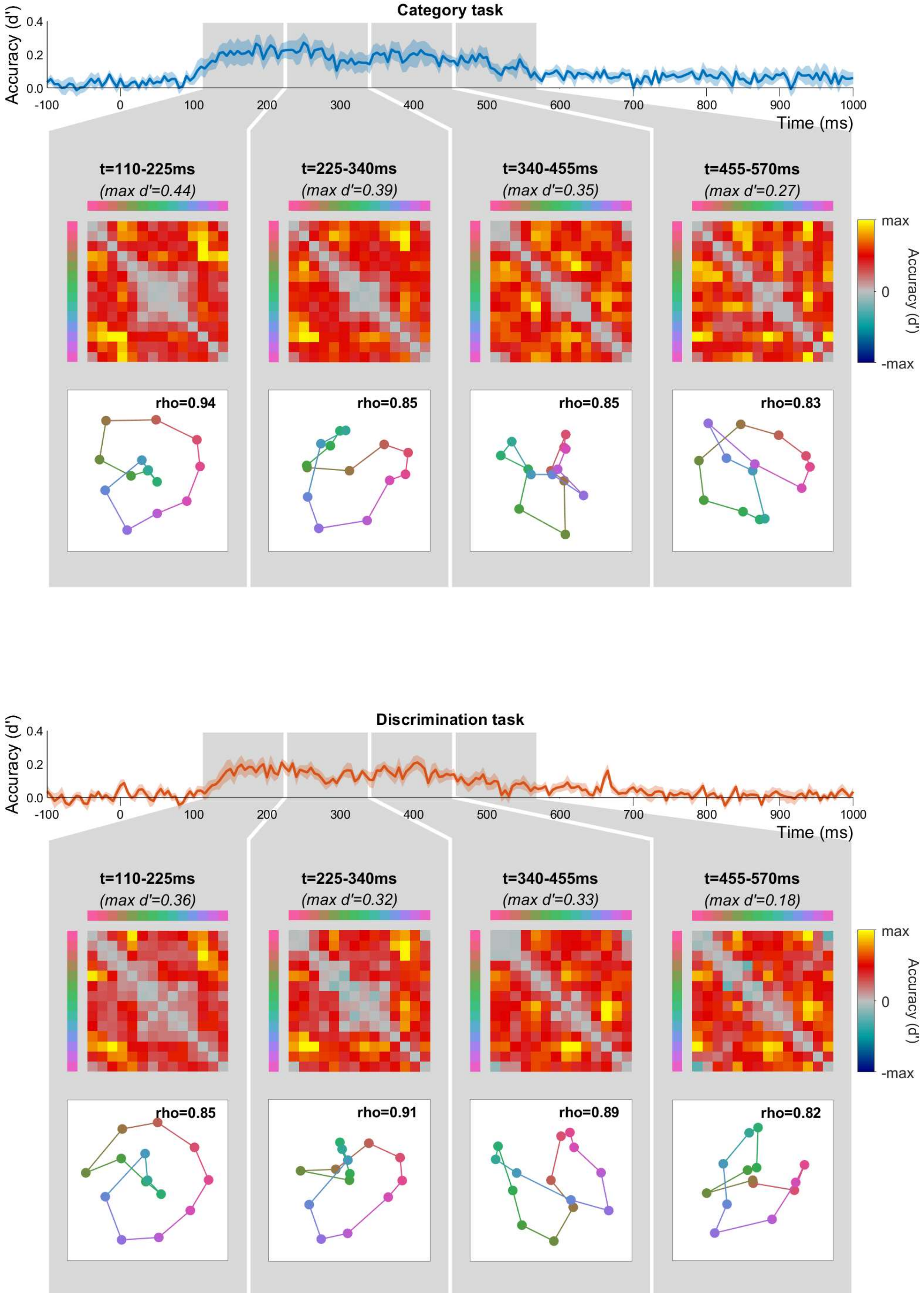


***Figure A6***: *Average dissimilarity matrices (DSMs) and corresponding MDS solutions for data from the category task (upper) and discrimination task (lower) for classifiers trained on data from the DVC. Plotting conventions as in Figure 6.*

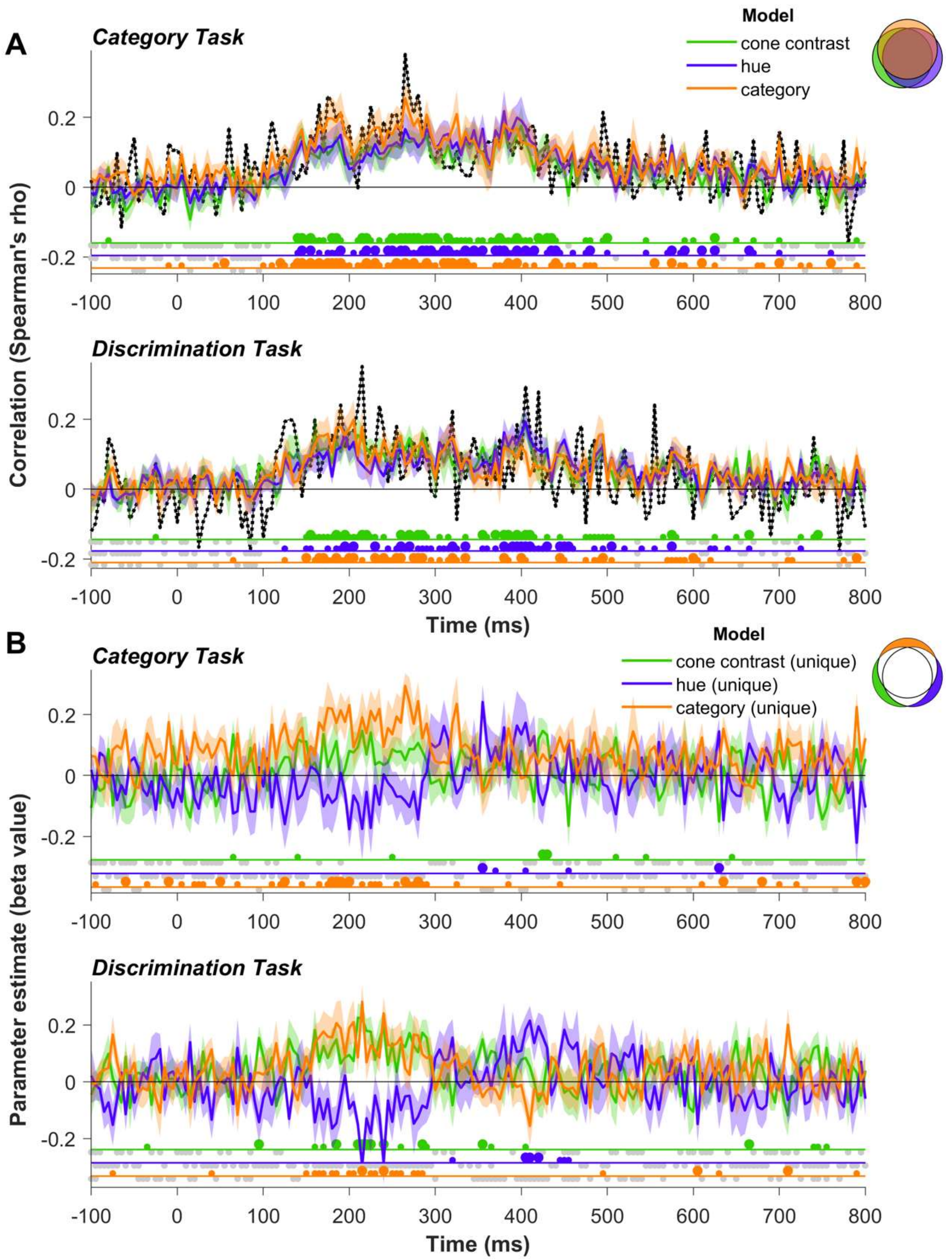


***Figure A7***: *Representational similarity analysis of hue decoding, for classifiers trained on data from the VVC. Plotting conventions as in Figure 7.*

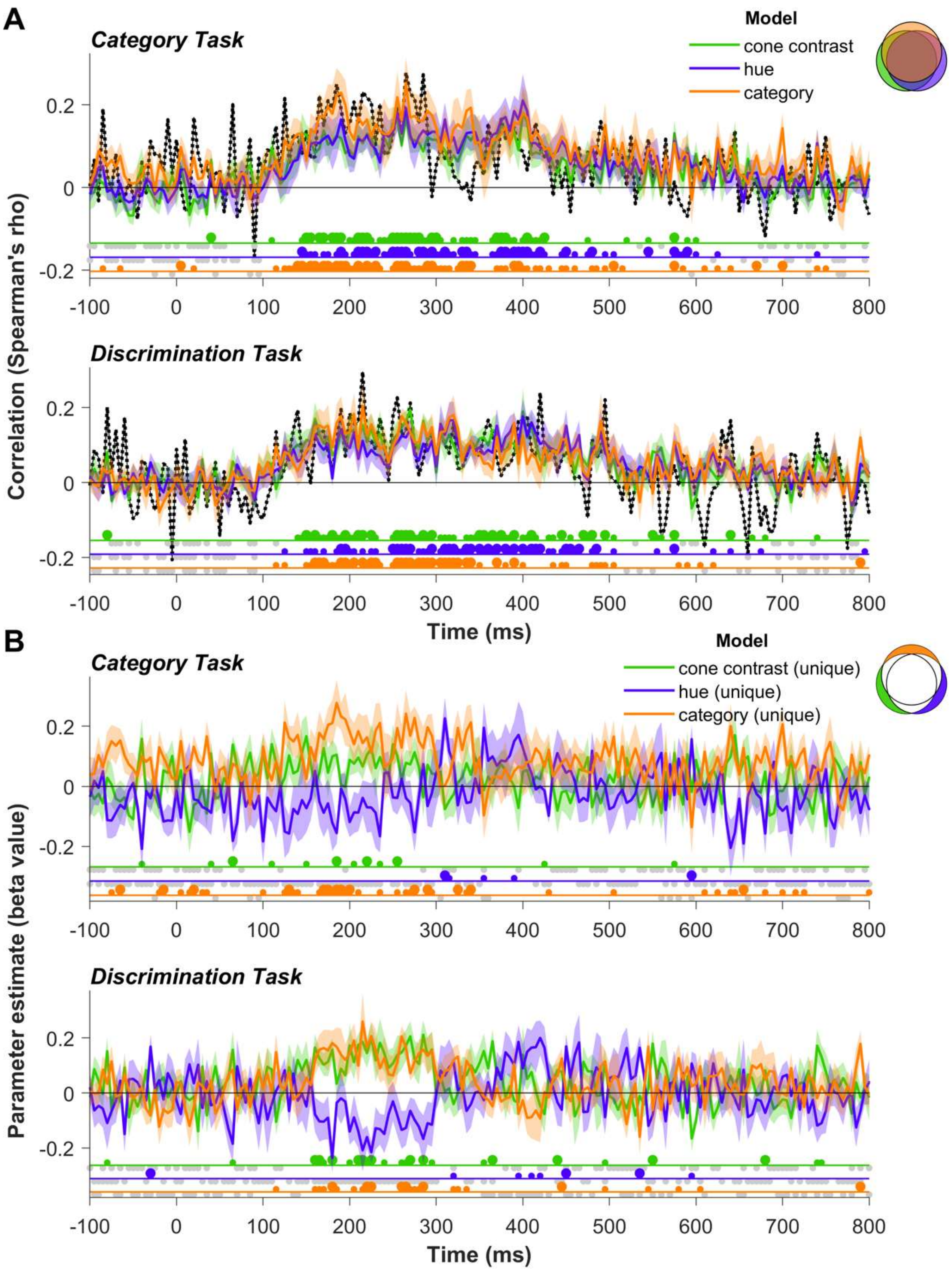


***Figure A8***: *Representational similarity analysis of hue decoding, for classifiers trained on data from the DVC. Plotting conventions as in Figure 7.*
